# A hybrid spectral library combining DIA-MS data and a targeted virtual library substantially deepens the proteome coverage

**DOI:** 10.1101/2020.01.16.909952

**Authors:** Ronghui Lou, Pan Tang, Kang Ding, Shanshan Li, Cuiping Tian, Yunxia Li, Suwen Zhao, Yaoyang Zhang, Wenqing Shui

## Abstract

Data-independent acquisition mass spectrometry (DIA-MS) is a rapidly evolving technique that enables relatively deep proteomic profiling with superior quantification reproducibility. DIA data mining predominantly relies on a spectral library of sufficient proteome coverage that, in most cases, is built on data-dependent acquisition-based analysis of the same sample. To expand the proteome coverage for a pre-determined protein family, we report herein on the construction of a hybrid spectral library that supplements a DIA experiment-derived library with a protein family-targeted virtual library predicted by deep learning. Leveraging this DIA hybrid library substantially deepens the coverage of three transmembrane protein families (G protein coupled receptors; ion channels; and transporters) in mouse brain tissues with increases in protein identification of 37-87%, and peptide identification of 58-161%. Moreover, of the 412 novel GPCR peptides exclusively identified with the DIA hybrid library strategy, 53.6% were validated as present in mouse brain tissues based on orthogonal experimental measurement.

## Introduction

Data-independent acquisition mass spectrometry (DIA-MS) is emerging as a powerful technology for proteomics research owing to its superior accuracy and reproducibility in proteomic quantification while retaining relatively deep coverage of the proteome (Gillet et al., 2012; Ludwig et al., 2018). To maximize the proteome coverage in DIA-MS analyses, sample-specific spectral libraries are typically built based on peptide identifications from a conventional data-dependent acquisition (DDA) experiment, which often involve offline pre-fractionation of peptide samples (Schubert et al., 2015). Raw DIA data can then be processed for peptide identification and quantification using a peptide-centric scoring algorithm (Ting et al., 2015) against this DDA experiment-derived spectral library. Alternatively, DIA data can be processed and searched directly against a FASTA sequence database (Ting et al., 2017; Tsou et al., 2015), however, such library-free approaches usually result in lower proteome coverage (Ting et al., 2017).

Recent innovative work by Gessulat *et al*. (Gessulat et al., 2019) and Tiwary *et al*. (Tiwary et al., 2019) demonstrated the feasibility of building a virtual spectral library based on separate predictions of fragment ion intensities and peptide retention times from deep learning models. Fundamentally, these major proof-of-concept studies demonstrated that DDA experiment-derived spectral libraries can be replaced with virtual spectral libraries built for experimentally-detected peptides to achieve nearly equivalent whole-proteome coverage (Gessulat et al., 2019; Tiwary et al., 2019).

In theory, current deep learning models can make predictions for all peptides yielded from *in silico* digestion of the entire proteome, but this would result in the substantial expansion of the spectral library and attendant significant increase of the false discovery rate (FDR) (Rosenberger et al., 2017). Given that many biological studies focus on a specific class of proteins, we propose a new strategy of constructing a targeted virtual library for a given protein superfamily to deepen its proteome coverage. Targeted MS assays have been developed for the detection and quantification of a predetermined set of proteins in complex matrices across multiple samples (Picotti and Aebersold, 2012). However, these conventional assays have several drawbacks. They require tremendous initial effort to select optimal peptides and establish assay parameters for each given protein and instrument platform. Also, only tens of proteins are routinely measured in a single run with targeted MS assays (Kusebauch et al., 2016). Herein, we present an approach of exploiting a targeted virtual library to profile the expression of hundreds of proteins within a superfamily by single-injection DIA analysis while strictly controlling the FDR in data mining.

## Results

### Generating an initial DIA spectral library for mouse brain tissues

To evaluate our strategy, we chose transmembrane protein families as our targets. These proteins were selected because of their strong hydrophobicity, relative low abundance, and fast turn-over, which make them challenging to profile using conventional proteomics techniques. Specifically, we focused on the G protein-coupled receptor (GPCR) superfamily: the mouse genome contains 524 annotated GPCRs, each having seven transmembrane domains (Katritch et al., 2013). Given that GPCRs represent one of the most prominent classes of drug targets, particularly for neurological diseases (Huang et al., 2017; Katritch et al., 2013), the ability to deeply and accurately profile the GPCR sub-proteome will greatly benefit both basic neuroscience and therapeutic development. However, GPCRs are notoriously under-represented in proteomic datasets. A meta-proteome analysis of diverse human samples identified only 65 GPCR proteins (Wilhelm et al., 2014) of the 831 encoded in the human genome, a much lower proportion than for most other protein superfamilies. A similarly low identification rate for GPCRs was also reported in a very deep proteomic survey of human cells (56 GPCRs identified) (Bekker-Jensen et al., 2017) (Supplementary Fig. 1).

Our study examined the GPCR sub-proteome in three mouse brain tissues: cerebellum, midbrain, and spinal cord. We prepared cell membrane fractions and performed single-injection DIA-MS analysis of digested protein extracts for each tissue, and identified 36, 38, and 38 GPCRs based on spectral matches for 405, 402, and 381 peptide precursors in the three brain regions, respectively (Supplementary Table 1). In our workflow, DIA data from 12 total analyses (from experimental quadruplets of each region) were directly searched against a mouse FASTA database using Spectronaut (Bruderer et al., 2015) to generate an initial spectral library (*i.e.*, without building a sample-specific spectral library from DDA experiments; Fig. 1). This DIA spectral library for the brain samples included 415 peptide precursors mapped to 38 GPCR proteins.

**Fig. 1.**
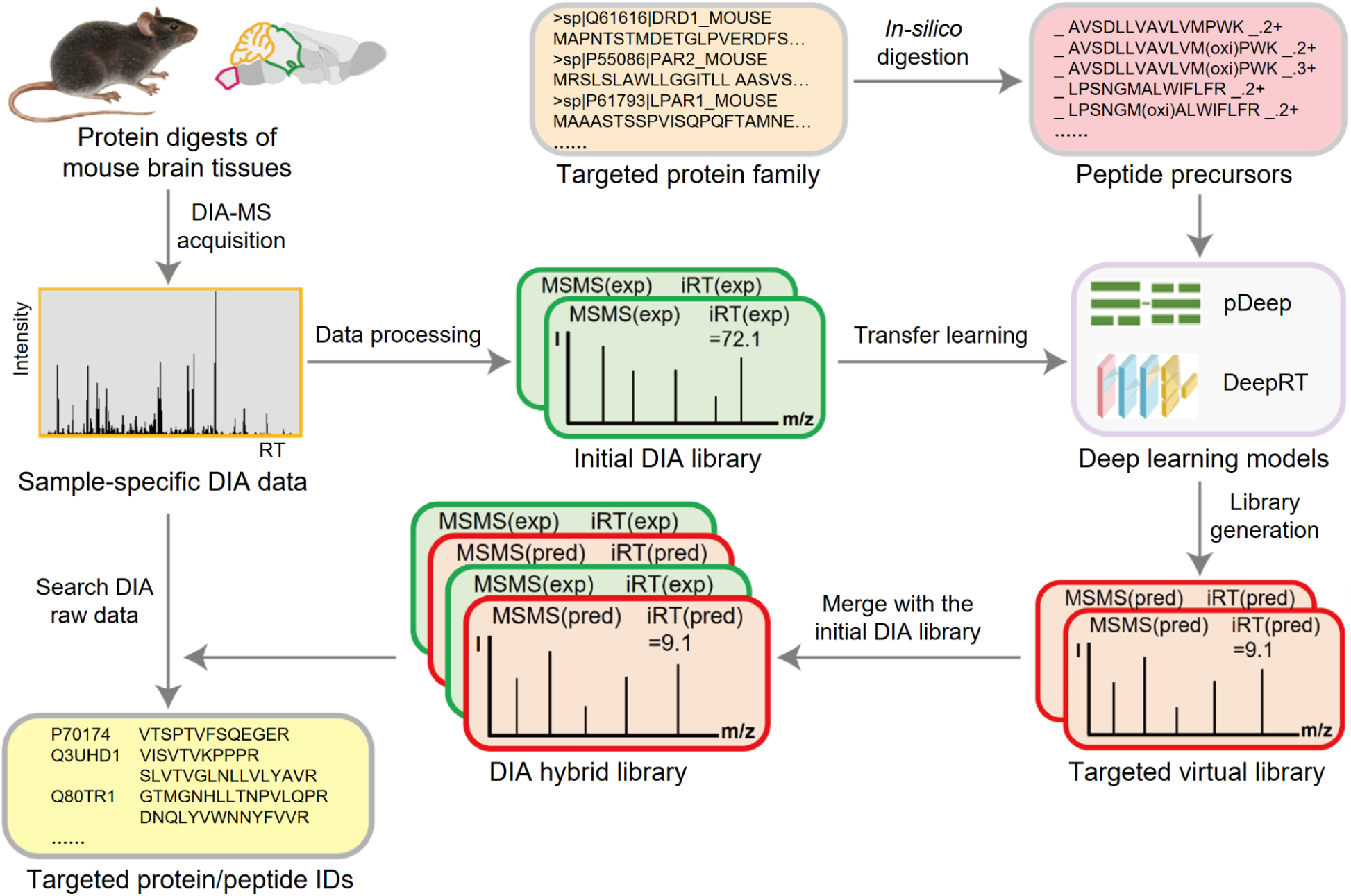
Overall workflow for DIA data mining using a hybrid spectral library. This hybrid library is constructed by merging a DIA experiment-derived library with a protein family-targeted virtual library built on all *in silico* digested peptide precursors. The targeted virtual library is generated using two deep learning models (pDeep and DeepRT) which are re-trained by transfer learning based on sample-specific DIA data alone. This study examines protein extracts of membrane fractions prepared from three mouse brain regions (cerebellum, midbrain, and spinal cord). ID, identification; pred, predicted; exp, experimental.

### Constructing a GPCR-targeted virtual library with re-trained pDeep and DeepRT

Before constructing a virtual spectral library, we tested the performance of several deep learning models to predict fragment ion intensities and retention time indices (iRT) for the 415 GPCR peptide precursors from the initial DIA spectral library. Distinct from the aforementioned whole-proteome virtual library approaches, we here used the deep neutral network-based models pDeep (Zhou et al., 2017) to predict fragment ion intensities and DeepRT (Ma et al., 2018) to predict iRT from GPCR peptide sequences (Fig. 1). The pre-trained pDeep model achieved excellent overall agreement between the experimental and predicted MSMS spectra at a defined collision energy for this GPCR peptide test set (median Pearson correlation coefficient = 0.93, median spectral angle = 0.85) (Supplementary Fig. 2A-B). However, for iRT prediction and because of large differences in the liquid chromatography conditions adopted in our DIA experiment vs. the earlier experiments upon which the DeepRT was originally trained (Ma et al., 2018), the initial performance of DeepRT was lower than expected (Supplementary Fig. 2C).

Unsatisfied with these pre-trained models, we next applied a transfer learning technique (Pan and Yang, 2010) to further train both the pDeep and DeepRT models using the majority of non-GPCR peptide entries in our DIA spectral library (27,390 in total). Subsequent analysis of our GPCR peptide test set showed significant improvements compared to the original pre-trained models (*e.g*., ΔiRT_95%_ value of 38 units for the pre-trained DeepRT vs. 12.9 units for the re-trained model; Supplementary Fig. 2). Notably, when we evaluated another deep learning model (Prosit) with our GPCR peptide test set, we found that the prediction results for both fragment ion intensities and iRTs were no better or slightly worse than those obtained with our re-trained pDeep and DeepRT models (Supplementary Fig. 3).

Having obtained sufficient prediction performance with the two deep learning models, we next set out to construct a GPCR-targeted virtual library. Specifically, we used the re-trained models to predict fragment ion intensities and iRT values for tryptic peptides yielded from an *in silico* digestion of the full complement of 524 GPCR proteins in the mouse genome (Fig. 1). To account for the known influence of digestion settings on the library size, we performed *in silico* digestion with 12 different combinations of peptide charge state, numbers of missed cleavages, and Met oxidation status (referred to as P1–P12; Fig. 2A).

**Fig. 2.**
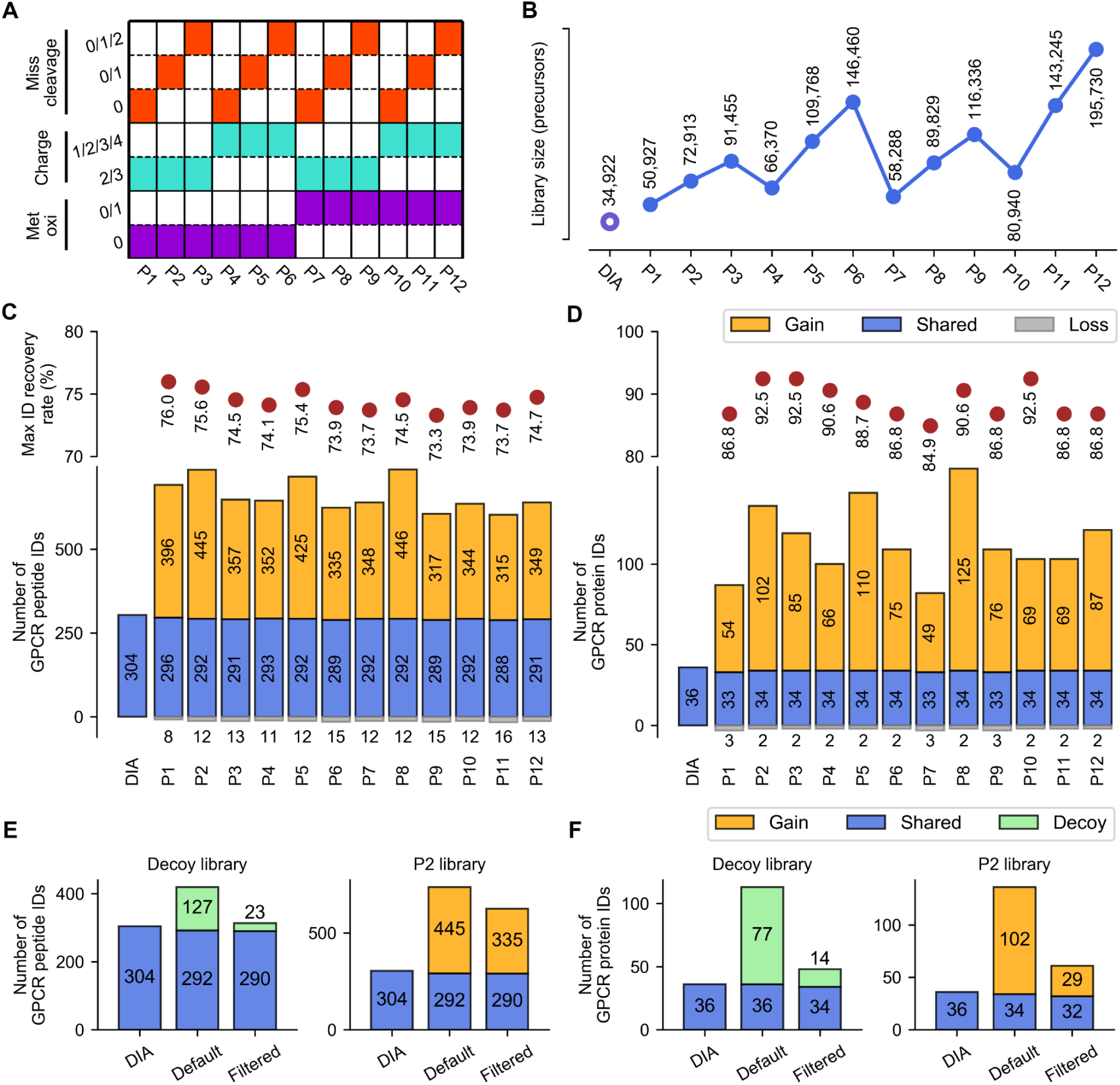
Increasing the depth of GPCR identification by DIA-MS with the hybrid library. **A**, *In silico* digestion of the 524 mouse GPCR proteins with 12 combinations (P1-P12) of peptide charge states, numbers of missed cleavages, and Met oxidation status. The peptide length is restricted to 7-33 residues. **B,** The number of peptide precursors in the initial DIA spectral library, and each “hybrid library” comprising the initial DIA spectral library plus the GPCR-targeted virtual library. **C-D,** Number of GPCR peptide identifications (IDs) (**C**) and protein IDs (**D**) in the cerebellum between the initial DIA library and 12 hybrid libraries. Relative to the number of peptide/protein IDs obtained with the initial DIA library (shown on the left), the proportion of shared IDs for each hybrid library is shown in blue, gained IDs in orange, and lost IDs in grey. The protein/peptide ID number in each fraction is annotated. Note that additional DDA experiments were conducted on pre-fractionated peptide samples from each brain region, and the GPCR identification lists from the initial DIA and new DDA experiments were merged to generate a “max ID list”. Max ID recovery rates refer to the percentages of *bona fide* identifications from these max ID lists, which were recovered using our 12 hybrid libraries. **E-F,** The number of GPCR peptide (**E**) or protein (**F**) IDs using the decoy hybrid library (left) or the P2 hybrid library (right) with default Spectronaut parameters (Default) or after data filtration based on Cscore >0.9 (Filtered). Relative to the protein/peptide IDs with the initial DIA library (left), the proportion of shared IDs is shown in blue, decoy IDs in green, and gained IDs in orange.

### Increasing the depth of GPCR identification by DIA-MS with a hybrid library

Each GPCR virtual library generated with a given set of conditions was then merged with the experimental DIA spectral library, yielding 12 distinct hybrid libraries. Whereas the initial DIA spectral library comprised 34,922 precursors, the size of the 12 hybrid libraries ranged between 50,927–195,730 precursors (Fig. 2B, Supplementary Table 2). Remarkably, searching our DIA data against each of the 12 hybrid libraries invariably led to drastically more putative GPCR identifications than searching with the initial DIA library. Taking the cerebellum sample as an example, an average of 114 GPCR proteins (from an average of 661 peptides) were putatively identified using the 12 virtual libraries, whereas only 36 proteins (from 304 peptides) were identified with the initial DIA library (Figs. 2C-D). In the best-performing library (P2), 737 peptides were mapped to 136 GPCRs, representing a gain of 445 peptides and 102 GPCRs relative to the initial DIA library. Of note, this prominent increase in coverage was only observed for GPCRs; that is, there was no significant change in the total numbers of protein identifications when using the 12 hybrid libraries (Supplementary Fig. 4, Supplementary Table 3). Moreover, similar increases in the numbers of putative GPCR peptides and proteins were also obtained when analyzing DIA data from the two other mouse brain regions (Supplementary Fig. 5).

### FDR assessment in use of the GPCR hybrid library

The huge gains in the number of putatively identified GPCRs prompted us to closely examine the potential for false positives since the expanded library size from the incorporation of a targeted virtual library was expected to increase the error rate (Rosenberger et al., 2017). We first performed an additional DDA experiment by fractionating the peptide samples from each brain region (Supplementary Table 4). Our concatenation of the GPCR identification lists from the initial DIA and the new DDA experiments yielded a “max identification (ID) list” (*i.e.*, the maximal collection of experimentally identified GPCR proteins and peptides). We then calculated the percentages of *bona fide* identifications from these max ID lists that were recovered using our 12 hybrid libraries. The P2 hybrid library yielded the highest recovery rates: 92.5% of proteins and 75.6% of peptides in the cerebellum sample (Figs. 2C-D), with similar results for the other two brain regions (Supplementary Fig. 5). Thus, P2 was chosen as the best-performing condition for virtual library construction (missed cleavages 0/1, charge state 2/3, Met oxidation 0). Furthermore, we observed no loss in the reproducibility of quantification for total peptides or for GPCR peptides identified with the P2 hybrid library compared to the initial DIA spectral library (median CVs at 8.90%-10.50%; Supplementary Fig. 6)

Beyond these *bona fide* identifications, searching with the P2 hybrid library putatively identified 372 novel GPCR peptides representing 87 novel GPCRs, which were not present in the cerebellum max ID list (Supplementary Fig. 7). To estimate the error rate amongst these P2-hybrid-library-only GPCR identifications, we created a decoy virtual library via the *in silico* digestion of 524 reverse GPCR sequences (again using the P2 condition). After combining this decoy virtual library with the initial DIA spectral library, a DIA data search with this hybrid library yielded 127 decoy peptides and 77 decoy proteins in the cerebellum (Figs. 2E-F; Supplementary Table 5). This result strongly indicated a significant FDR when using the default settings in Spectronaut, although we set FDR to <1% at the peptide and protein levels in the DIA data search. When we subsequently applied a more stringent data filtration cut-off (Cscore >0.9), 81.9% of the decoy peptides and 81.8% of the decoy proteins were removed, and the same data filtration retained 625 (84.8%) GPCR peptides and 61 (44.9%) GPCRs identified in the cerebellum with the P2 hybrid library (Figs. 2E-F). The other two regions showed similar results before and after data filtration (Supplementary Fig. 8, Supplementary Table 5).

Altogether 71 GPCR proteins and 810 GPCR peptides were identified from three mouse brain regions after applying the data filtration cut-off, compared to 38 GPCR proteins and 310 peptides identified with the initial DIA library (Figs. 4A, 4D). Thus, with an appropriately controlled error rate by data filtration, the use of our targeted hybrid library to process DIA data can substantially deepen the coverage for a given protein superfamily beyond conventional proteomic data acquisition and processing techniques.

**Fig 4.**
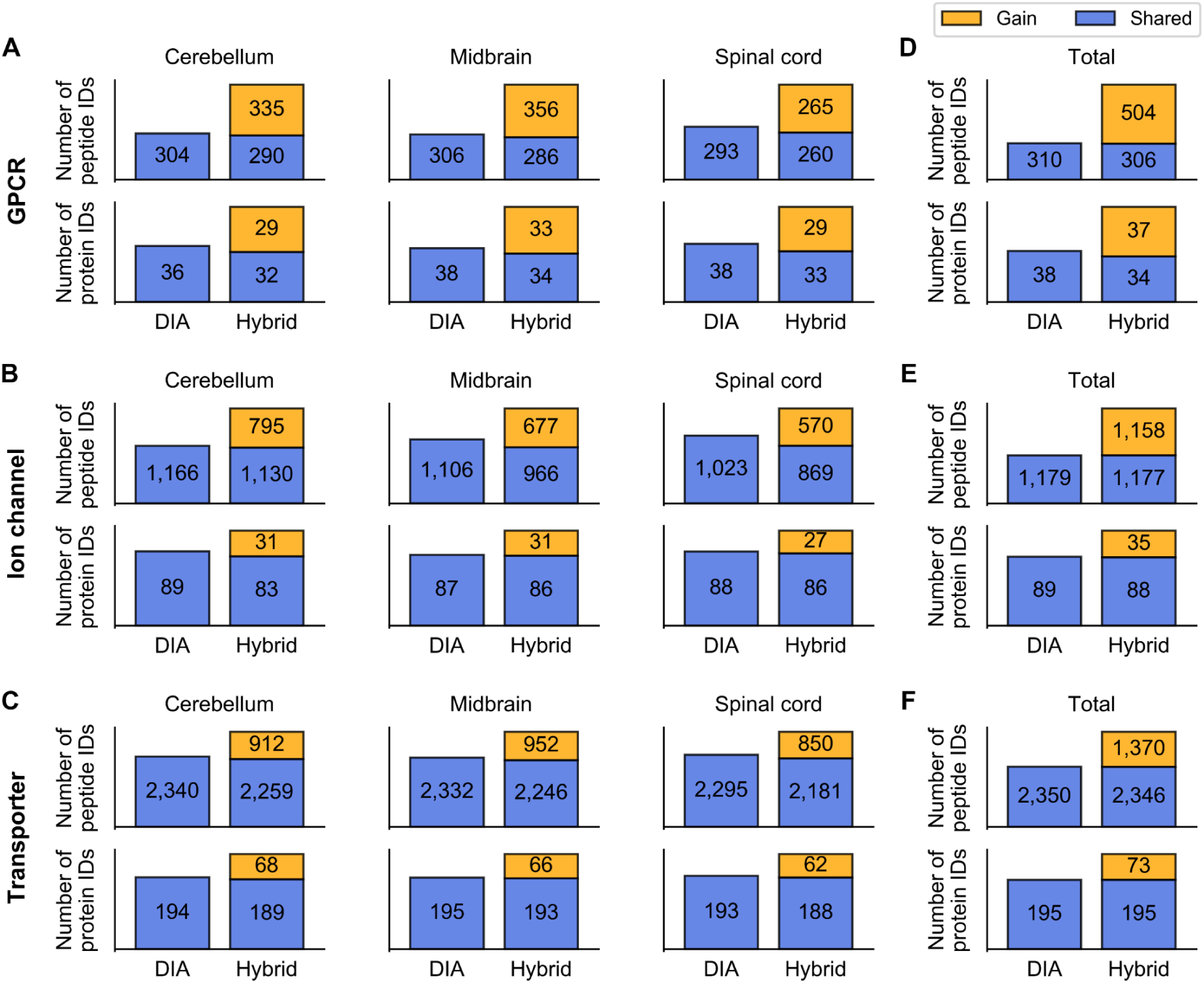
Comparison of GPCR, ion channel and transporter identifications (IDs) in mouse brain tissues using the initial DIA library (with default software settings) and the hybrid library P2 (after data filtration). **A-C**, Comparison of GPCR (A), ion channel (B), and transporter (C) peptide IDs (upper) and protein IDs (lower) in three brain regions. **D-F**, Comparison of total non-redundant GPCR (D), ion channel (E), and transporter (F) peptide IDs (upper) and protein IDs (lower) identified from three regions. In each panel, relative to the protein/peptide IDs with the initial DIA library (shown on the left), the proportion of shared IDs for hybrid library P2 is shown in blue, and gained IDs in orange. The number of protein IDs in each fraction is annotated.

### Validation of novel GPCR peptides exclusively identified with the hybrid library

Compared to the max ID lists, there were 412 non-redundant novel GPCR peptides identified in three mouse brain regions after data filtration (Supplementary Table 6). These peptides represent novel identifications beyond any DIA- or DDA-based experimental measurement and they were only obtained using our DIA hybrid library strategy. Next, we designed experiments to empirically validate their existence in the protein digests of the specific brain regions (Fig. 3A). We first conducted targeted MS assays on novel peptides in each region by parallel-reaction-monitoring (PRM) analyses. High-quality MSMS spectra were obtained for 214 peptides (Supplementary Table 7). When we incorporated these experimental fragment ion intensities and measured iRTs into the initial DIA spectral library, searching the DIA data with Spectronaut default parameters verified the identification of 207 novel peptides in the mouse brain tissues (Supplementary Table 7). Thus, 50.2% of peptides identified exclusively using our DIA hybrid library approach are indeed present in the samples. Given that individual biological replicates were prepared for the DIA and PRM experiments, we speculate that the invalidated portion may result from relatively low reproducibility of extracting and analyzing these novel peptides most of which are at low abundance.

**Fig. 3.**
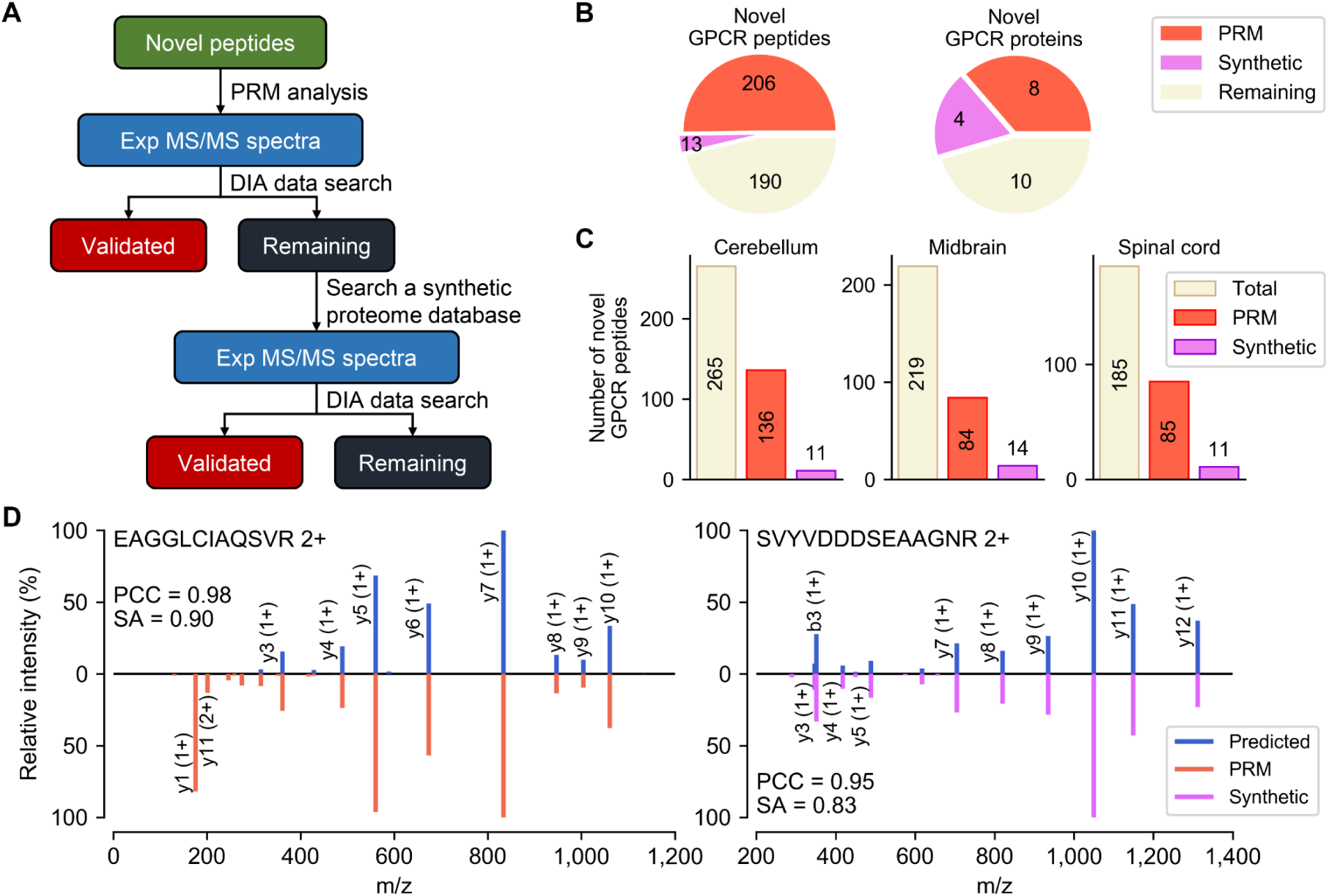
Validation of GPCR peptides exclusively identified with the DIA hybrid library strategy. **A**, Workflow of novel peptide validation, based on targeted MS (PRM) analysis or on searching the synthetic human proteome database in ProteomeTools. Novel GPCR peptides were validated through DIA data searching of experimental MSMS spectra acquired from the PRM experiment or the synthetic peptide spectral repository. **B,** Total number of novel GPCR proteins or peptides identified in the three brain regions, validated based on the PRM experiment (red) or the synthetic peptide spectral repository (pink). Also shown are the remaining unconfirmed peptides (beige). **C,** Total number of novel GPCR peptides identified in each brain region using our DIA hybrid library strategy (beige), as well as the number validated based on the PRM experiment (red) or the synthetic peptide spectral repository (pink). **D,** Pseudo mirror plots of example MSMS spectra comparing the predicted (upper) and experimental (lower) fragmentation patterns for two novel GPCR peptides. The experimental MSMS spectrum was acquired from the PRM experiment for peptide EAGGLCIAQSVR (left) or from the synthetic reference spectral repository for peptide SVYVDDDSEAAGNR (right). The Pearson correlation coefficient (PCC) and normalized spectral contrast angle (SA) are indicated. PRM, parallel reaction monitoring.

Still pursuing validation of the remaining 205 putative novel GPCR peptides, we searched for synthetic peptides of the same sequences deposited to the ProteomeTools spectral libraries (Zolg et al., 2017). The MSMS spectra retrieved for the 23 synthetic peptides were incorporated into the initial DIA spectral library for DIA data searching, which resulted in the validation of 22 additional novel GPCR peptides (Supplementary Table 7). Thus, a total of 221 non-redundant novel peptides were validated with experimental MSMS data (Fig. 3B), corresponding to a total validation rate of 53.6% (Supplementary Table 6). In detail, the GPCR peptide validation rates achieved with our DIA hybrid library strategy in the three brain regions are 55.5%, 44.8%, and 51.9%, respectively (Fig. 3C). These validated novel peptides confirmed the existence of 12 out of 22 novel GPCR proteins only discovered with the DIA hybrid library (Fig. 3B). Mirror plots for two example peptides comparing the predicted fragmentation patterns alongside experimental MSMS spectra from either the PRM analysis or the synthetic human proteome database reflect strong agreement between prediction and measurement. (Figs. 3D-3E).

### Deepening the proteome coverage for other transmembrane protein families or for GPCRs using a published dataset

To extend our strategy to deepen coverage for transmembrane proteins of other families, we employed the same workflow described in Fig. 1 to re-train the deep learning models with sample-specific DIA data and to construct hybrid spectral libraries (under the P2 optimal condition) for the 240 ion channels and the 452 transporters encoded by the mouse genome (Supplementary Fig. 9). These protein family-targeted virtual libraries (used in concert with the stringent data filtration) enabled a substantial increase of the proteome coverage for both ion channels (from 89 proteins and 1,179 peptides identified in the three regions with the initial DIA library to 123 proteins and 2,335 peptides with the hybrid library) and transporters (from 195 proteins and 2,350 peptides identified with the initial DIA library to 268 proteins and 3,716 peptides with the hybrid library) (Figs. 4B, 4C, 4E, 4F; Supplementary Table 8).

To avoid re-training models using different datasets for different protein families, we built generic models of pDeep and DeepRT with 90% of randomly selected precursors from the initial DIA library to construct virtual libraries for three transmembrane protein families (Supplementary Fig. 10). The generic models showed very similar performance to the protein family-specific models. Furthermore, DIA hybrid libraries generated based on prediction by the generic models yielded peptide identifications for three transmembrane protein families very comparable to the family-specific models (Supplementary Fig. 10). Therefore, it is feasible to build generic models from an experimentally acquired DIA dataset to predict for any selected protein family.

Finally, seeking to demonstrate the effectiveness of our strategy for mining previously acquired DIA data from other sources, we downloaded a published DIA dataset that was acquired from total lysates of the brain barrel cortex regions from newborn mice (Bruderer et al., 2017). We used the same procedure for model re-training and processing of the DIA dataset obtained at 4 time points each in technical triplicate (12 analyses in total) (Supplementary Fig. 11). Re-trained models were implemented to construct a targeted virtual library for all mouse GPCRs with the P2 condition, which was then incorporated with the initial spectral library generated from the DIA dataset to yield a hybrid library. Compared to 27 GPCR proteins and 229 peptides in total identified with DIA data alone over the time course, our DIA hybrid library strategy enabled exceptionally deep mapping of 83 GPCRs based on spectral matches of 895 peptides after data filtration (Fig. 5, Supplementary Table 9).

**Fig. 5.**
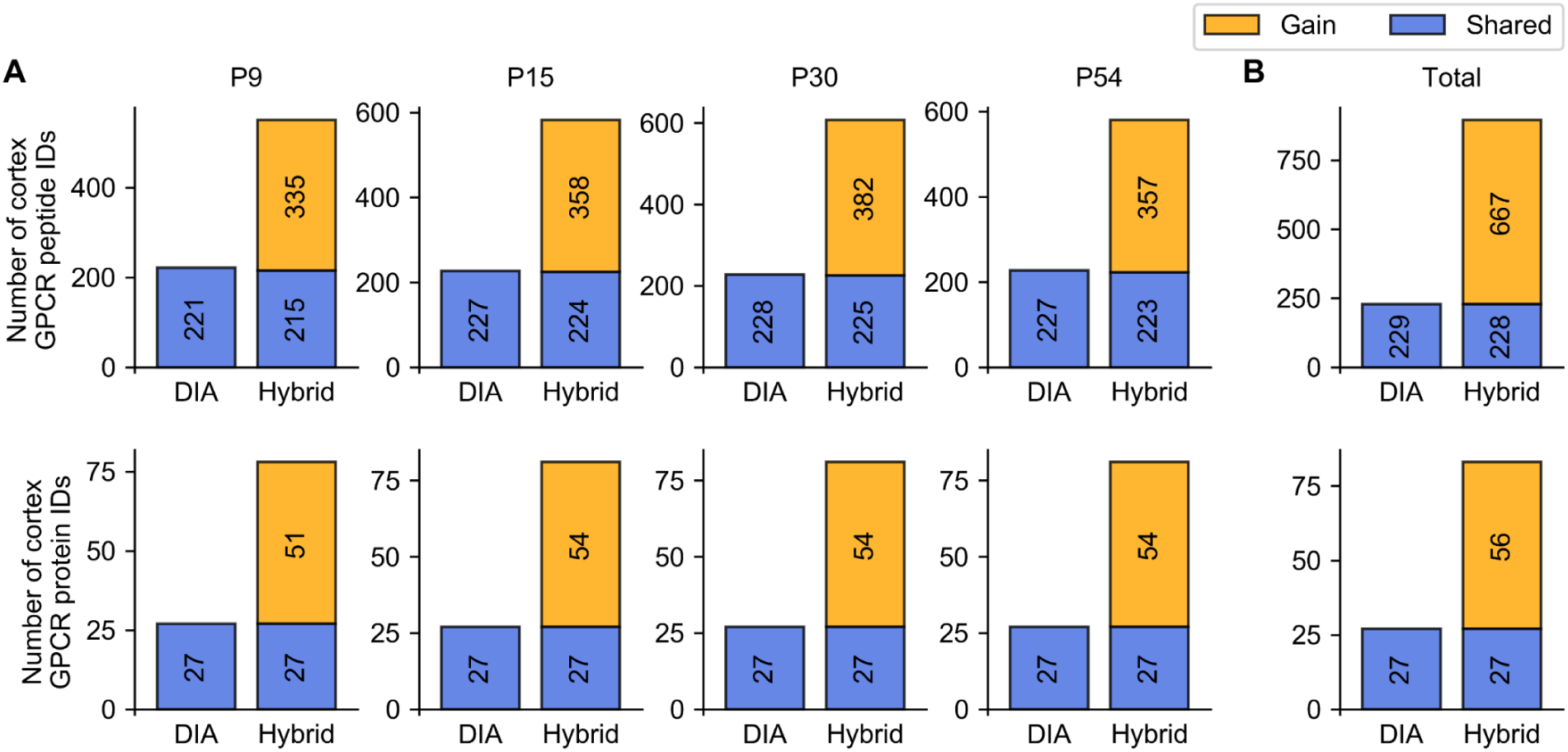
DIA hybrid library substantially increases GPCR-targeted proteome coverage in the analysis of a published dataset. **A**, Comparison of GPCR peptide (upper) and protein (lower) IDs in mouse brain cortex at four time points (P9, P15, P30, P54) during development. **B,** Comparison of the total non-redundant GPCR peptide (upper) and protein (lower) IDs identified in this time course. Relative to the number of protein/peptide IDs with the initial DIA library (with default software parameters; left), the proportion of shared IDs for the P2 hybrid library (after data filtration; right) is shown in blue and gained IDs in orange. The number of protein IDs in each fraction is annotated.

## Discussion

Compared to previous studies that demonstrated the applicability of deep learning-based models to both DDA and DIA data analysis (Gessulat et al., 2019; Tiwary et al., 2019), our study explores a fundamentally new direction for DIA data mining. In previous work, a virtual spectral library was first generated for all experimentally detected peptides from a sample-specific spectral library derived from DDA analysis. Subsequent replacement of the entire experimental spectral library with the virtual library achieved a whole-proteome coverage close to the experimental library. In contrast, our study aims to deepen the sub-proteome coverage for a selected protein family that surpasses experimental limits. We constructed a hybrid spectral library that supplemented a DIA experiment-derived library with a protein family-targeted virtual library built on all *in silico* digested peptide precursors.

Our study demonstrates how DIA data mining using this hybrid library strategy can remarkably increase the depth of proteomic profiling for a selected protein family without compromising FDR control or quantification reproducibility. As a proof-of-concept, GPCR identifications in three mouse brain regions were increased from 310 peptides mapped to 38 proteins with the initial DIA library (using default software parameters) to 810 peptides mapped to 71 proteins with the DIA hybrid library (after stringent data filtration). Moreover, 412 novel GPCR peptides and 22 novel GPCR proteins were not observed in any conventional mining of data from DIA or pre-fractionation-based DDA experiments. Importantly, we performed orthogonal PRM experiments and exploited the synthetic human proteome database to validate the existence of 53.6% novel GPCR peptides detected exclusively with our DIA hybrid library strategy.

It is important to emphasize that our workflow for deep learning-based virtual library construction and subsequent generation of hybrid libraries only requires a small DIA dataset, which can be acquired from minimal sample quantities and minimal instrument time. Thus, our DIA hybrid library approach circumvents the need for the time-consuming generation of sample-specific spectral libraries through extensive pre-fractionation of peptide samples. Furthermore, this strategy is adaptable to targeting multiple protein families and to previously acquired DIA datasets, allowing for retrospective data mining based on new hypotheses. Although not directly showcased here, we anticipate that this bioinformatics strategy will enhance the detection of members in any protein family (or any set of selected proteins) of biological interest, including splicing isoforms, sequence variants, or proteins bearing specific PTMs, to an unprecedented depth.

### Limitations of the Study

Our current study deepens the sub-proteome coverage for three transmembrane protein families based on the DIA-MS dataset acquired from mouse brain tissues. It remains to be investigated whether we can exploit this DIA hybrid library strategy to map other protein families from the same dataset in largely increased depth. Moreover, as we need to construct virtual libraries for individual pre-determined protein families, it would be more efficient to combine them and build a larger hybrid library for DIA-MS data search. But we also recognize the challenge of FDR control with an expanded hybrid library. Thus it is necessary to further develop the decoy library method for accurate assessment of FDR in DIA-MS data search with a hybrid spectral library.

## Methods

### Animals and brain tissue preparation

Animals were used in accordance with standard ethical guidelines. All experimental procedures were approved by the Institutional Animal Care and Use Committee of ShanghaiTech University, China. Adult C57BL/6J male mice (8-10 weeks old at the time of surgery) were bred in-house and kept in a temperature- and humidity-controlled environment under a 12 h light / dark cycle with food and water available ad libitum. The mice were euthanized with 2% chloral hydrate. Each brain was rapidly removed from the skull and manually dissected to obtain three different regions (cerebellum, midbrain, and spinal cord) under the microscope. The brain regions were rinsed in ice-cold PBS buffer, then transferred into individual tubes, immediately weighed, frozen on LN2, and stored at −80 °C until analysis.

### Membrane protein extraction and digestion

The membrane fraction of each mouse brain region was prepared according to a previously reported procedure (Suski et al., 2014). Briefly, the same brain region dissected from 6-7 mice was pooled and homogenized in the buffer of 300 mM sucrose, 0.5% BSA, 100 mM EDTA, 30 mM Tris/HCl, pH 7.4 supplemented with protease inhibitor (Roche). Crude membrane fractions were isolated from the homogenate by ultra-centrifugation at 160,000 g at 4°C for 1 h. The membrane pellet was solubilized in 4% SDS and 100 mM DTT in 100 mM Tris/HCl, pH 7.6, and heated at 95°C for 5 min for full denaturation and disulfide breakage. Protein concentration was determined using the BCA assay (TIANGEN, Beijing, China).

Digestion of membrane proteins was performed using the FASP method described previously (Wisniewski et al., 2009). Briefly, about 20 μg of protein extract from each region was diluted in 8 M urea, 50 mM NH4HCO3 and exchanged to the same buffer using the 30 KDa MWCO centrifugal filter unit (Satorious, Germany) by centrifugation at 13,000 g at 4°C for 20 min. The following centrifugation steps were performed under the same condition. Subsequently, 100 μl of 50 mM iodoacetamide in 8 M urea, 50 mM NH4HCO3 was added to the concentrate and incubated at room temperature in darkness for 30 min prior to centrifugation. The concentrate was diluted with 200 μl 50 mM NH4HCO3 and centrifuged again, and this step was repeated twice. Proteins were digested with sequencing-grade trypsin (Promega, Madison, USA) at an enzyme-to-protein ratio of 1:100 (w/w) at 37°C for 3 h, followed by the addition of trypsin at 1:50 (w/w) and incubation at 37°C overnight. After acidification, the protein digest was desalted with C18-SepPak columns (Waters, Milford, USA) and lyophilized under vacuum. For each brain region, protein digestion and peptide desalting was performed in quadruplicate. Half of the peptide sample from each replicate was spiked in with the iRT reference kit (Biognosys, Zurich, Switzerland) and saved for DIA analysis, the other half from the same region was pooled for offline fractionation and DDA analysis.

### Offline fractionation of protein digests

For each brain region, the proteolytic digest pooled from quadruplicate preparation (40 μg) was loaded onto an equilibrated, high-pH, reversed-phase fractionation spin column (Thermo Fisher Scientific). Peptides were bound to the hydrophobic resin and desalted by washing the column with water. A step gradient of increasing acetonitrile concentrations in a volatile high-pH elution solution was then applied to elute bound peptides into 8 different fractions which were collected by centrifugation. All these peptide fractions were then dried in a vacuum centrifuge and stored until mass spectrometry analysis.

### Mass spectrometry analysis

#### DIA/DDA data acquisition

The nano LC-MS/MS analysis was conducted on an EASY-nLC 1000 connected to Orbitrap Fusion Tribrid mass spectrometer (Thermo Fisher Scientific, USA) with a nano-electrospray ionization source. Peptides were separated on an analytical column (200 mm x 75 μm) in-house packed with C18-AQ 3 μm C18 resin (Dr. Maisch, GmbH, Germany) over a 120-min gradient from 5% to 35% mobile phase B (0.1% FA in acetonitrile) at a flow rate of 300 nl/min. For each brain region, the protein digests from four experimental replicates were injected separately for DIA analysis. In addition, the offline peptide fractions of protein digests from each region were injected for DDA analysis. In the DIA mode, the resolution of Orbitrap analyzer was 60,000 for MS1 and 30,000 for MS2. 35 variable windows were set to cover the mass range of 350-1250 *m/z.* The automatic gain control (AGC) target was set to 1e6 in MS1 and 5e5 in MS2, with a maximum ion injection time of 20 ms in MS1 and 50 ms in MS2. In the DDA mode, MS1 and MS2 resolution were the same as in DIA with a precursor mass range of 300-1700 *m/z*, and the isolation window was 1.6 m/z. The AGC target was set to 4e5 in MS1 and 1e5 in MS2, with a maximum ion injection time of 50 ms in MS1 and 50 ms in MS2. Top 12 peptide precursors were automatically selected for MS2 data acquisition, with a collision energy at 30%±5%.

#### PRM data acquisition

In the PRM experiment, the membrane protein extract of each brain region was re-prepared from fresh mouse brain tissue, and underwent tryptic proteolysis according to the previous procedure. Peptide samples were analyzed by Q-Exactive HF mass spectrometer coupled to EASY-nLC 1200 (Thermo Fisher Scientific). Peptides were eluted on the same analytical column over the same LC gradient as described for DIA/DDA analysis. The precursor inclusion lists were created based on the identifications of novel GPCR peptides in each region (exclusively identified with DIA hybrid library and absent in the max ID list). Each inclusion list contained the precursor information for 60-80 novel GPCR peptides. In the PRM mode, the resolution of Orbitrap analyzer was 60,000 for MS1 and 30,000 for MS2. The AGC target was set to 3e6 in MS1 and 5e5 in MS2, with a maximum ion injection time of 30 ms in MS1 and 100 ms in MS2. The isolation window was set to 1 m/z, and collision energy at 30%. For each brain region, the protein digest was injected in a duplicate set to screen all novel GPCR peptide precursors split up into 4-5 inclusion lists. For the remaining un-verified precursors, a second protein digest was prepared and injected in a duplicate set to screen all remaining GPCR peptide precursors split up into 2-3 inclusion lists.

### Mass spectrometry data processing

#### DIA spectral library generation

This initial DIA library was generated by searching all DIA raw data acquired from 3 brain regions (12 runs in total) against mouse SwissProt/Uniprot FASTA database (state 2019/05/21, containing 17015 entries) plus the iRT-kit FASTA format sequences using the Spectronaut^TM^ 12.0 software (Biognosys, Zurich, Switzerland). Standard settings were adopted as follows: Trypsin/P as the specific protease; maximal two missed cleavages per peptide allowed; carbamidomethyl (C) as fixed modification; oxidation (M) and acetylation (at protein N-terminus) as variable modifications; false discovery rates (FDR) on PSM/peptide/protein levels all set to 1%. The resulting DIA spectral library contained 34922 precursor entries corresponding to 3149 protein groups. This initial DIA spectral library was used for subsequent DIA data searching, model re-training, and testing, as well as hybrid library construction.

#### DIA data search

The 12 DIA raw data were analyzed using Spectronaut^TM^ 12.0 in the library-dependent mode. Either the initial DIA library or a hybrid library for a specific protein family was imported. Spectronaut processing was performed with default settings: minimum relative intensity 5%, best six (minimum 6) fragment ions per spectrum, automatic iRT calibration, XIC RT width automatic, MS1 and MS2 mass tolerance dynamic, decoy limit dynamic, decoy generation method ‘mutated’, the protein and precursor Q-value cutoff both at 0.01. Standard reports for all searches were generated to collect the unique peptide and protein entries under each condition. Cscore and other parameters are all exported to the searching result file (.xls).

#### DDA data search

For each brain region, the eight DDA raw data files acquired from peptide fractionations were combined and searched against the mouse SwissProt/Uniprot FASTA database (state 2019/05/21) supplemented with a contaminant database using Proteome Discoverer^TM^ search engine (Version 2.2, Thermo Fisher Scientific) with the following settings: Trypsin/P as the specific protease; maximal two missed cleavages per peptide allowed; carbamidomethyl (C) as fixed modification; oxidation (M) and acetylation (at protein N-terminus) as dynamic modifications; mass tolerance for precursor ions was set to 10 ppm and for fragment ions to 0.02 Dalton. False discovery rates (FDR) on PSM/protein/peptide levels were all set to 1%. For each brain region, the GPCR protein and peptide identifications (IDs) from either DIA data search or DDA data search were merged to yield the max ID list containing non-redundant protein/peptide IDs.

#### PRM data search

The PRM raw data files obtained from each brain region were combined and searched against the mouse SwissProt/Uniprot FASTA database (state 2019/05/21) supplemented with a contaminant database using Proteome Discoverer^TM^ search engine (Version 2.2, Thermo Fisher Scientific) with the same settings in DDA data search. Any GPCR peptides identified with identical sequence, modification and charge state to those in the PRM inclusion list were retained, and their MSMS spectra were manually inspected to confirm peptide identifications (⩾6 fragment ions and each ion intensity >10^4) The fragmentation data for these novel GPCR peptides were exported for further construction of a hybrid library (described below).

### Re-training and testing models

#### Data preprocessing

In the initial DIA library, peptide precursors and the corresponding fragmentation ion entries were selected as the training set if they meet the following criteria: peptide length of 7-26 residues, charge state of +1 to +5, at least six fragment ions assigned in the PSM, not mapped to any mouse GPCR protein sequences. According to the distribution of all peptide and fragment ion coordinates in the initial DIA library (Supplementary Fig. 12), precursors in this training set represent the majority (>95%) in the initial DIA library and they have high-quality MSMS spectra. The MSMS and iRT data of 27390 non-GPCR precursors in the training set were used for re-training models through transfer learning. The initial DIA library also contained 412 GPCR peptide precursors and their MSMS/iRT data served as a test set for model evaluation. To re-train and test models for other targeted protein families (ion channels and transporters), we re-filtered precursors in the initial DIA library to obtain the training set, and the subset of precursors mapped to the targeted family members were selected as the test set.

#### pDeep model re-training and testing

pDeep2 is downloaded from github.com/pFindStudio/pDeep (Commit 2019/03/11). The training set was first transformed to the pLabel format as specified in the software package. Transfer learning was applied to the pre-trained model *pretrain-180921-modloss* using the script transfer_train.py with certain parameters modified: min_var_mod_num is set to 0; max_var_mod_num to 3; NCE to 0.3; instrument type to QE. Both the pre-trained and re-trained models were implemented using the script predict.py to predict fragmentation patterns for peptide precursors in the test set. Model prediction was performed with varying epoch values (5, 10, 15, 20, 30, 40, 50, 75, 100). Model performance was evaluated based on the MSMS spectrum similarity between predicted and experimental results (described below). The best pDeep model with an optimal epoch was obtained separately for each transmembrane protein superfamily.

#### MSMS spectrum similarity calculation

MSMS spectrum similarity analysis reflects the similarity of fragment ion distribution and intensities between two PSMs for the same precursor. Fragment ion intensities in the predicted or experimental spectrum are base-peak normalized. Only matched fragments between the predicted and experimental spectrum are retained for calculation. For example, if the predicted and experimental spectrum contains 6 and 7 fragment ions respectively and 5 are shared in two spectra, the m/z and intensity values for the 5 shared fragments are used for similarity calculation. We employed two methods to evaluate MSMS spectrum similarity. Pearson correlation coefficient (PCC) is calculated with the stats.pearsonr function in python package Scipy, which is defined as:

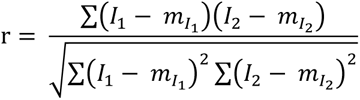

where *m*_*I*_1__ is the mean of vector *I*_1_ and *m*_*I*_2__ is the mean of vector *I*_2_.

Normalized spectral contrast angle is calculated as follows:

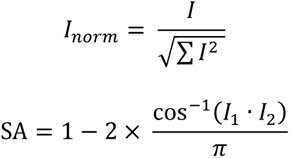

where *I* is the vector of intensity and *I_norm_* is the normalized intensity vector. *I*_1_ and *I*_2_ are two vectors of normalized intensity. A model that yields higher median PCC and SA gives better performance.

#### DeepRT model re-training and testing

DeepRT is downloaded from github.com/horsepurve/DeepRTplus (Commit 2018/11/25). The training set was first transformed to the format as specified in the software package. The model parameters were modified to be compatible with our experimental conditions as follows: min rt is set to −80; max rt to 176; time scale to 1 (*i.e.* in minute); max length to 66. Transfer learning was applied to the pre-trained model dia_all_epo20_dim24_conv8_filled.pt using the script capsule_network_emb.py. Both the pre-trained and re-trained models were implemented using the script prediction_emb.py to predict RTnorm values for peptides in the test set. iRT was then converted from RTnorm using the following equation (maximal and minimal RT were pre-defined in the model):

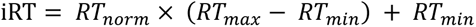

Model prediction was performed with varying epoch values (5, 10, 15, 20, 30, 40, 50, 75, 100). Model performance was evaluated using two parameters: ΔiRT95% (reflecting the deviation of predicted data from experimental data for 95% data points) and correlation coefficient of linear regression R^2^ (reflecting the correlation of predicted and experimental iRT values) which is calculated as below:

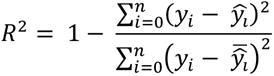

where 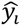 is the predicted retention time by regression function y = x. A model that yields smaller ΔiRT95% and larger R^2^ gives better performance. The best DeepRT model with an optimal epoch was obtained separately for each transmembrane protein family.

#### Prosit model testing

We implemented Prosit as an online server on the ProteomeTools website (https://www.proteomicsdb.org/prosit/) for prediction. As Prosit does not allow prediction of peptides with more than 30 residues, we transformed GPCR peptides within 30 residues in the test set to the Prosit input format, and collision energy was set to 30 (consistent with our acquisition method). After submitting to Prosit, predicted fragmentation and iRT data were compared with experimental data in the test set for model performance evaluation as described above.

### *In silico* digestion of a targeted protein family

For the GPCR family, protein sequences of all 524 members in the mouse proteome were subjected to *in silico* digestion under 12 different combination of conditions. These conditions vary in the number of missed cleavages (0, 0/1 or 0/1/2), Met oxidation (0 or 0/1), and charge state (2/3 or 1/2/3/4) of the yielded peptides. The maximal range of each digestion parameter covers >99% of all precursors present in the initial DIA library (Supplementary Fig. 12). Peptide length was set to be 7 to 33 residues, which also covers >99% precursors in the initial DIA library. For ion channel and transporter protein families, *in silico* digestion of all family members was conducted under the optimal condition P2. We also performed *in silico* digestion of 524 decoy GPCRs in reversed sequences under condition P2.

### Construction of virtual and hybrid libraries

Re-trained models of pDeep and DeepRT were implemented to predict fragment ion intensities and iRT values for all peptides yielded from *in silico* digestion of a targeted protein superfamily under a specific set of conditions. Then a virtual spectral library was built for a targeted superfamily based on the predicted results using an in-house script. This virtual library was merged with the initial DIA library to generate a hybrid library. All peptides assigned to the targeted protein superfamily in the initial DIA library were kept in the hybrid library. Peptides not identified in the DIA experiment yet predicted by our models were added to the initial DIA library. The resulting hybrid library for each protein superfamily was then used to search DIA raw data with default settings in Spectronaut 12 as described above. For the search results with a hybrid library, we filtered the protein and peptide identification reports based on Cscore >0.9.

A GPCR decoy library was generated based on prediction for all decoy peptides yielded from *in silico* digestion of decoy GPCRs as described above. This decoy library was also merged with the initial DIA library to generate a decoy hybrid library which was used to search DIA raw data. The resulting search results were filtered based on Cscore >0.9.

To validate the identifications of novel GPCR peptides, we also built two types of hybrid spectral libraries. One was generated by merging the initial DIA library with fragmentation data for 214 novel GPCR peptides that were identified in the PRM experiment. For any novel GPCR peptide repeatedly identified in PRM analysis, its best experimental MSMS spectrum was selected manually. The other hybrid library was generated by merging the initial DIA library with fragmentation data for 23 novel GPCR peptides that were retrieved from the synthetic peptide spectral repository in ProteomeTools (FTMS_HCD_30_annotated_190725.zip). These two hybrid libraries based on experimental MSMS spectra were then used to search DIA raw data with default settings in Spectronaut 12 as described above.

### Processing a published DIA dataset with the virtual library

The barrel cortex DIA dataset provided by Kelstrup *et al*. zx is downloaded from ProteomeXChange with identifier PXD005573. It contains 12 raw DIA data files acquired at 4 time points. This DIA dataset was imported to Spectronaut 12 to generate an initial DIA spectral library as described above. The resulting DIA library contains a total of 109611 peptide precursors mapped to 5153 proteins. It was filtered using the aforementioned criteria to retain 55242 non-GPCR peptide precursors with high-quality MSMS spectra for subsequent model re-training by transfer learning as described above. The subset of GPCR peptides in the initial DIA library served as the test set for evaluating the model performance before and after transfer learning (Supplementary Fig. 10). Next, *in silico* digestion was performed on 524 mouse GPCR proteins under condition P2. Fragment ion intensities and iRT for yielded peptides were predicted using the re-trained models of pDeep and DeepRT, which gave rise to a GPCR virtual library. This virtual library was merged with the initial DIA library to create a hybrid library. Finally, the raw DIA dataset was searched with Spectronaut 12 against the initial DIA library or the hybrid library so as to compare the number of GPCR protein/peptide identifications. For the search results with a hybrid library, we filtered the protein and peptide identification reports based on Cscore >0.9.

## Acknowledgement

We very much thank the fruitful discussion with Prof. Simin He and Dr. Wenfeng Zeng from Institute of Computing Technology, CAS, and their help with using pDeep 2. This work was funded by ShanghaiTech University, the National Program on Key Basic Research Project of China (2018YFA0507004 (W.S.), 2016YFA0501900 (Y.Z.), 2016YFA0501904 (Y.Z.), 2016YFC0905900 (S.Z.), 2018YFA0507000 (S.Z.)), and National Natural Science Foundation of China (31971362, 31671428, 31971178).

## Author contribution

R.L. and K.D. performed model re-training, library generation and MS data processing. P.T. prepared samples, acquired and analyzed PRM data. S.L. prepared samples and acquired DIA data. C.T. and Y.L. helped with brain sample preparation and MS data acquisition. S.Z. and Y.Z. were involved in the overall project management. W.S., R.L. and P.T. wrote the manuscript with edits from all authors. W.S. conceived and supervised the project.

## Supplementary Information

### Contents

Supplementary Figure 1. GPCR identifications in previous proteomic studies.

Supplementary Figure 2. Evaluation of pDeep and DeepRT model performance.

Supplementary Figure 3. Evaluation of Prosit model performance.

Supplementary Figure 4. Comparison of total protein identifications (IDs) in each brain region using the initial DIA library or different DIA hybrid libraries.

Supplementary Figure 5. Comparison of GPCR identifications (IDs) in two other brain regions using the initial DIA library and different hybrid libraries.

Supplementary Figure 6. Reproducibility of total peptide and GPCR peptide quantification across four replicates using the initial DIA library and the hybrid library P2.

Supplementary Figure 7. Comparison of GPCR identifications (IDs) in the max ID list of each brain region with those obtained using different hybrid libraries.

Supplementary Figure 8. Data filtration removed a large portion of decoy GPCR peptides in two other brain regions.

Supplementary Figure 9. Testing re-trained models of pDeep and DeepRT in prediction for members in ion channel and transporter families.

Supplementary Figure 10. Building generic models that can predict for all three transmembrane protein families.

Supplementary Figure 11. Testing models of pDeep and DeepRT re-trained with the published barrel cortex DIA dataset.

Supplementary Figure 12. Properties of stripped peptides or precursors in the initial DIA library.

Note: Supplementary Tables 1-9 are provided as separate Excel sheets.

**Supplementary Figure 1.**
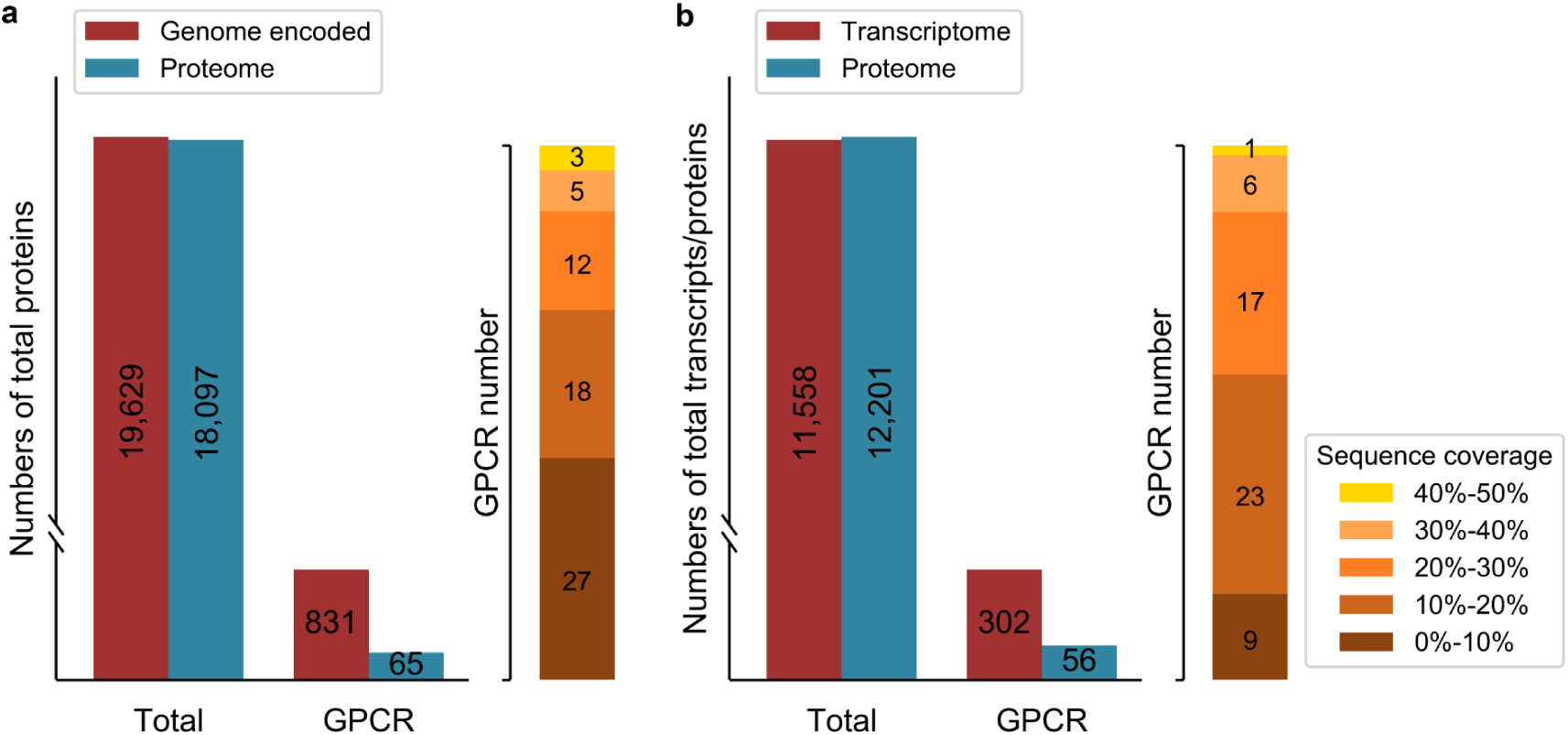
GPCR identifications in previous proteomic studies. (a) Numbers of total proteins and GPCR proteins reported in a metaproteome dataset from the analysis of various human tissues, cell lines and body fluids^1^ (blue bars) in comparison to numbers of all protein products and GPCRs encoded in the human genome (red bars). (b) Numbers of total proteins and GPCR proteins reported in a very deep proteomic survey of HeLa cells^2^ (blue bars) in comparison to numbers of all transcripts and GPCR transcripts detected in HeLa cells by RNA-seq^3^ (red bars). Orange bars designate the number of GPCRs in each category with the sequence coverage in a specific range. The majority of GPCRs identified in these two studies (57 out of 65 in a, 49 out of 56 in b) show sequence coverages below 30%.

---

**Supplementary Figure 2.**
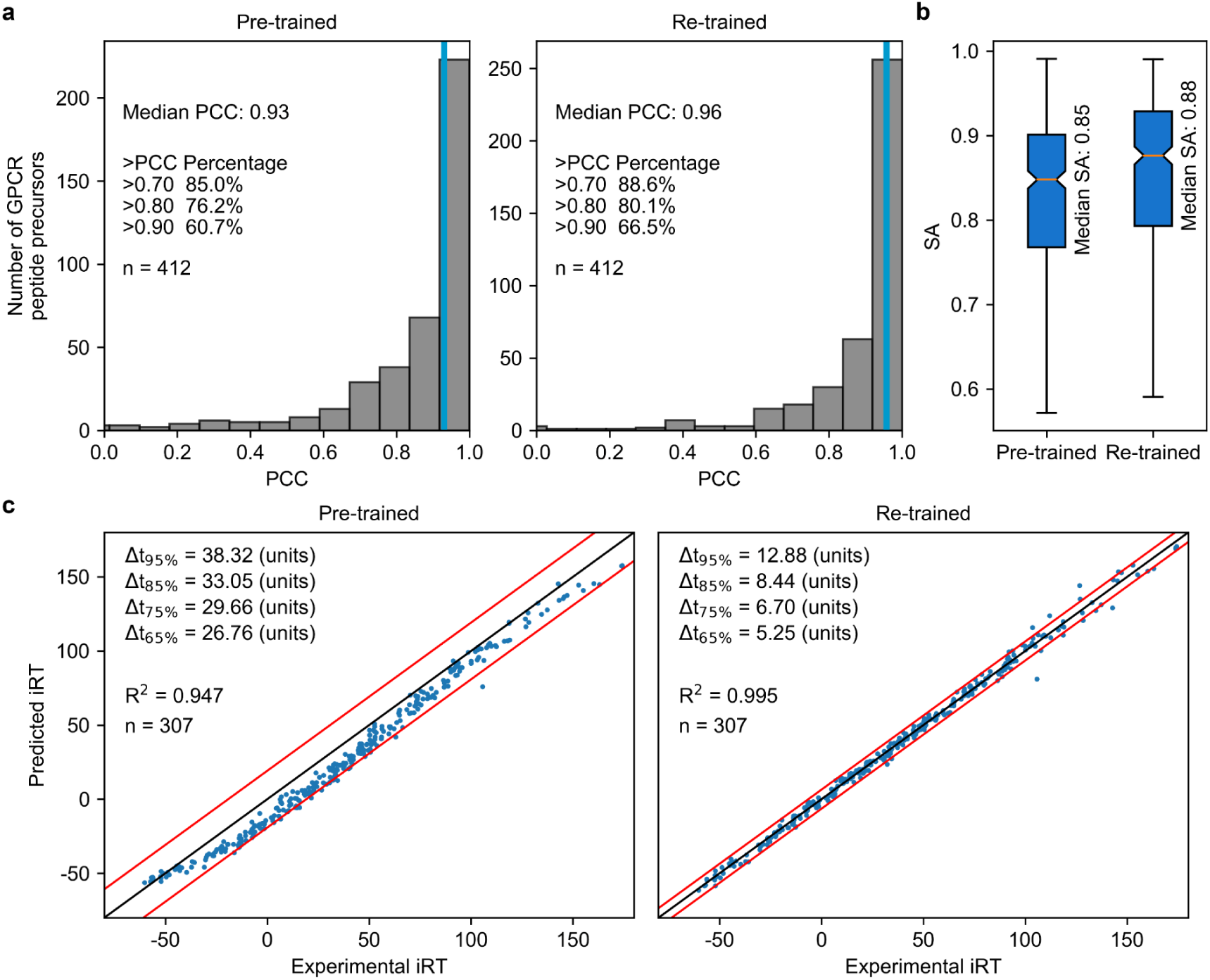
Evaluation of pDeep and DeepRT model performance. (a) Evaluation of pre-trained and re-trained models of pDeep based on distribution of Pearson Correlation Coefficient (PCC) calculated between predicted and experimental MSMS spectra. Vertical blue lines indicate the median PCC in the histogram; n is the number of GPCR peptide precursors in the test set. (b) Evaluation of pre-trained and re-trained models of pDeep based on the distribution of spectral contrast angle (SA) calculated between predicted and experimental MSMS spectra. Median SA is indicated. (c) Evaluation of pre-trained and re-trained models of DeepRT based on correlation of predicted and experimental iRT values. Correlation coefficient of linear regression (R^2^) is indicated; n is the number of GPCR peptides in the test set. Red lines mark the iRT window required to encompass 95% of all peptides (Δ_t95%_) around the diagonal (black line).

**Supplementary Figure 3.**
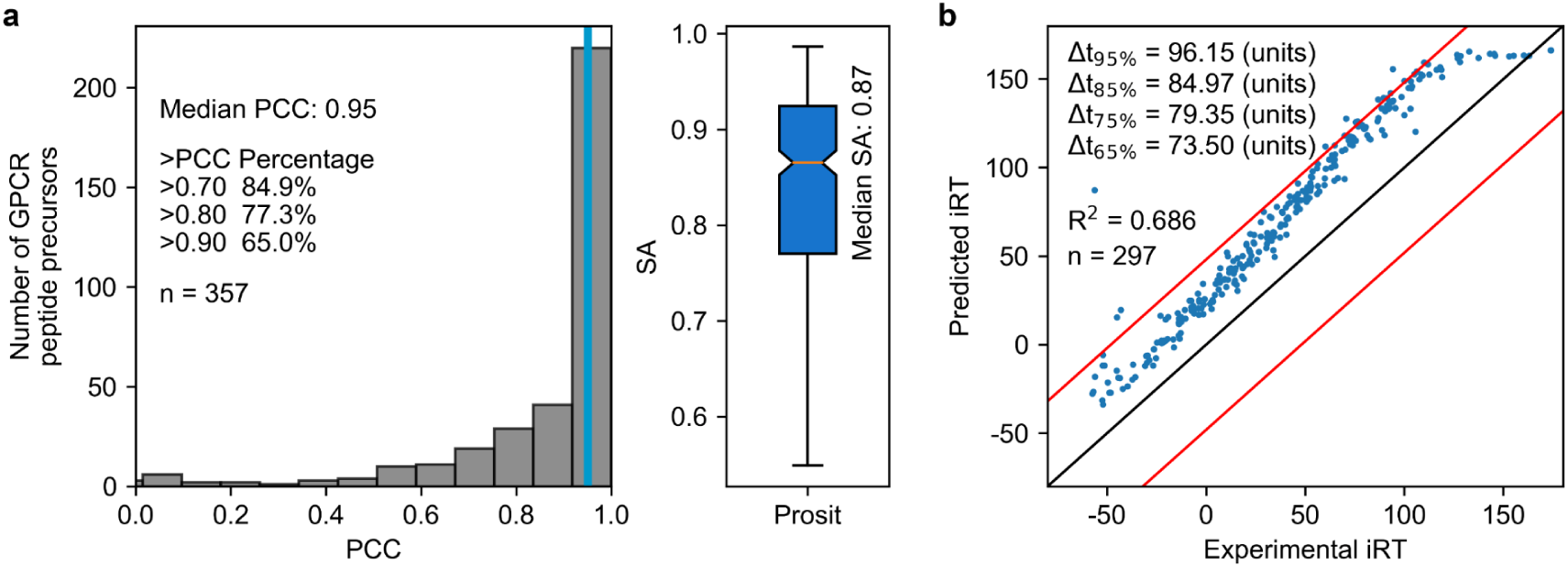
Evaluation of Prosit model performance. (a) Evaluation of fragmentation pattern prediction with the model based on distribution of PCC (left) and SA (right) calculated between predicted and experimental MSMS spectra. (b) Evaluation of iRT prediction with the model based on correlation of predicted and experimental iRT values. Correlation coefficient of linear regression (R^2^) is indicated. Red lines mark the iRT window required to encompass 95% of all peptides (Δ_t95%_) around the diagonal (black line). The number of GPCR peptide precursors or peptides with no more than 30 residues in the test set (n) is indicated in each panel.

**Supplementary Figure 4.**
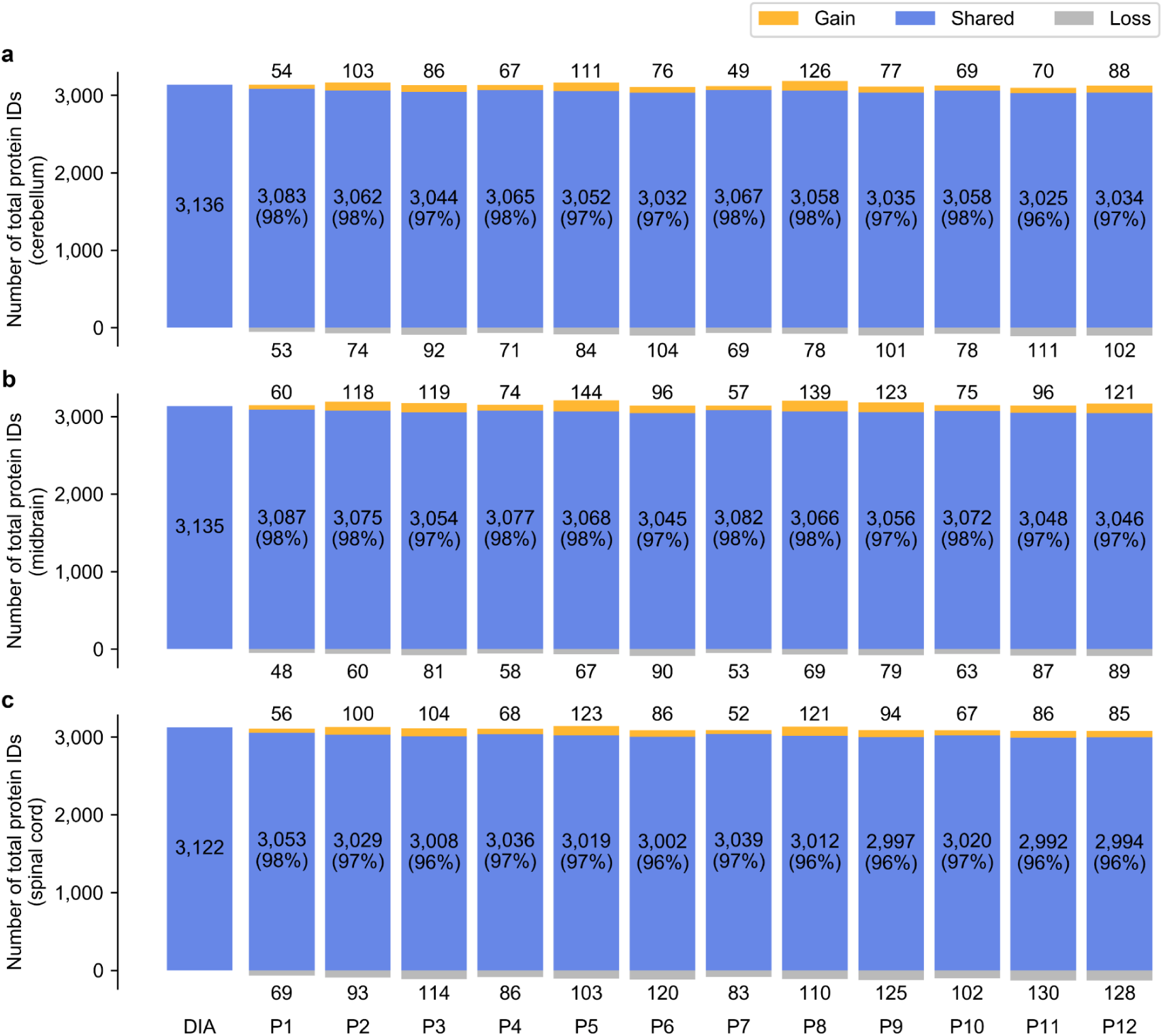
Comparison of total protein identifications (IDs) in brain regions of cerebellum (a), midbrain (b), and spinal cord (c) using the initial DIA library or different DIA hybrid libraries. Relative to the total protein IDs obtained with the initial DIA library (shown on the left), the proportion of shared IDs for each hybrid library is shown in blue; gained IDs are in orange; lost IDs are in grey. The number of protein IDs in each fraction is annotated.

**Supplementary Figure 5.**
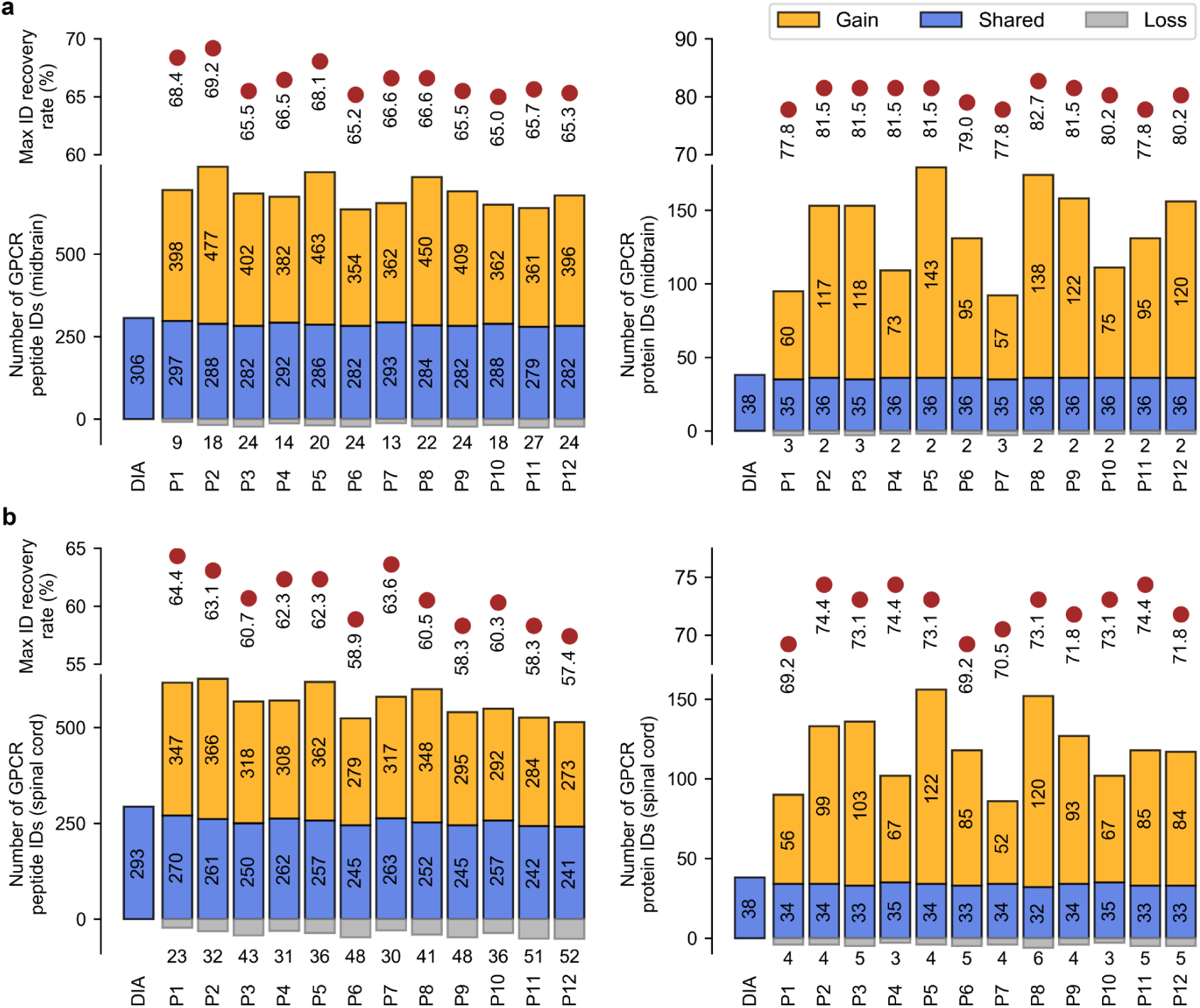
Comparison of GPCR identifications (IDs) in two other brain regions using the initial DIA library and different hybrid libraries. (a) Comparison of GPCR peptide IDs (left) and protein IDs (right) in midbrain. (b) Comparison of GPCR peptide IDs (left) and protein IDs (right) in spinal cord. Relative to the protein/peptide IDs with the initial DIA library (shown on the left), the proportion of shared IDs for each hybrid library is shown in blue; gained IDs are in orange; lost IDs are in grey. The number of protein IDs in each fraction is annotated. Max ID recovery rates refer to the percentages of *bona fide* identifications from respective max ID lists which were recovered using our 12 hybrid libraries (see details in Figure 2 legend).

**Supplementary Figure 6.**
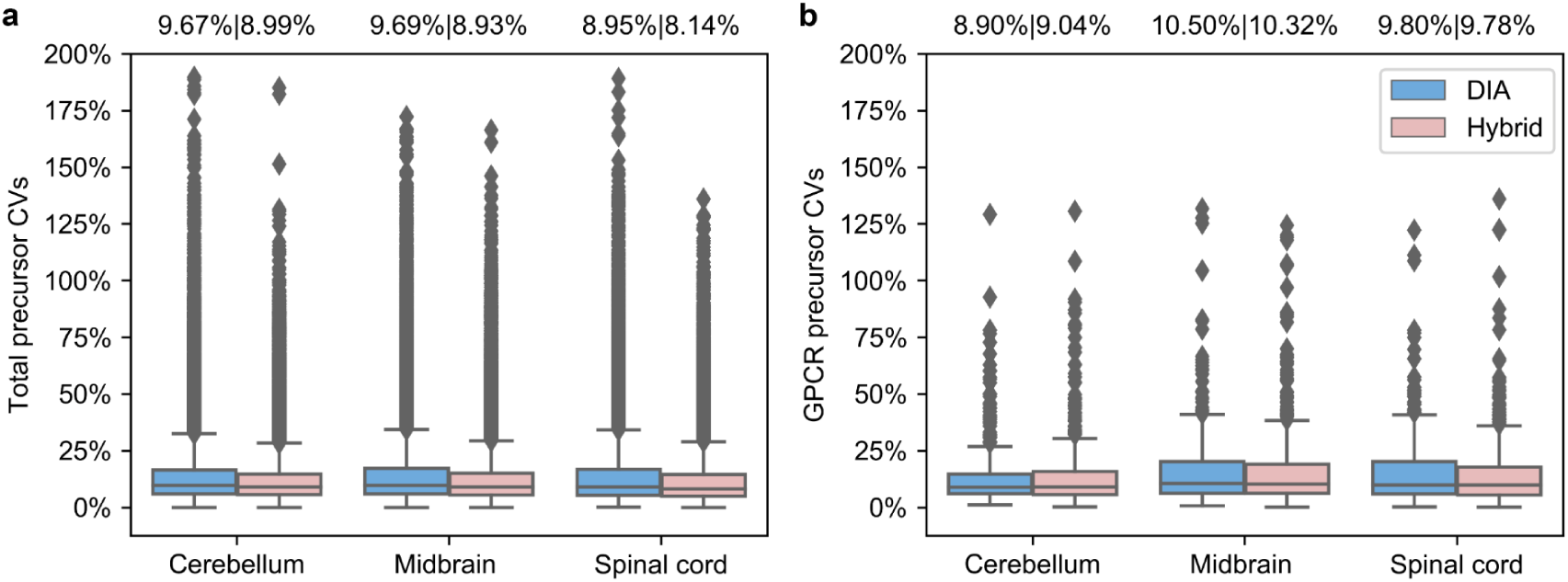
Reproducibility of total peptide and GPCR peptide quantification across four replicates using the initial DIA library and the hybrid library P2. Box plots of CVs for total peptide (a) and GPCR peptide (b) quantification using the initial DIA library (blue) and the hybrid library P2 (pink) are shown. Median CVs are annotated on top of each box plot. The lower and upper hinges correspond to the first and third quartiles (the 25th and 75th percentiles). The upper whisker extends from the hinge to the largest value no further than 1.5 × IQR (the inter-quartile range) from the hinge. The lower whisker extends from the hinge to the smallest value at most 1.5 × IQR from the hinge. Data beyond the end of the whiskers are plotted individually.

**Supplementary Figure 7.**
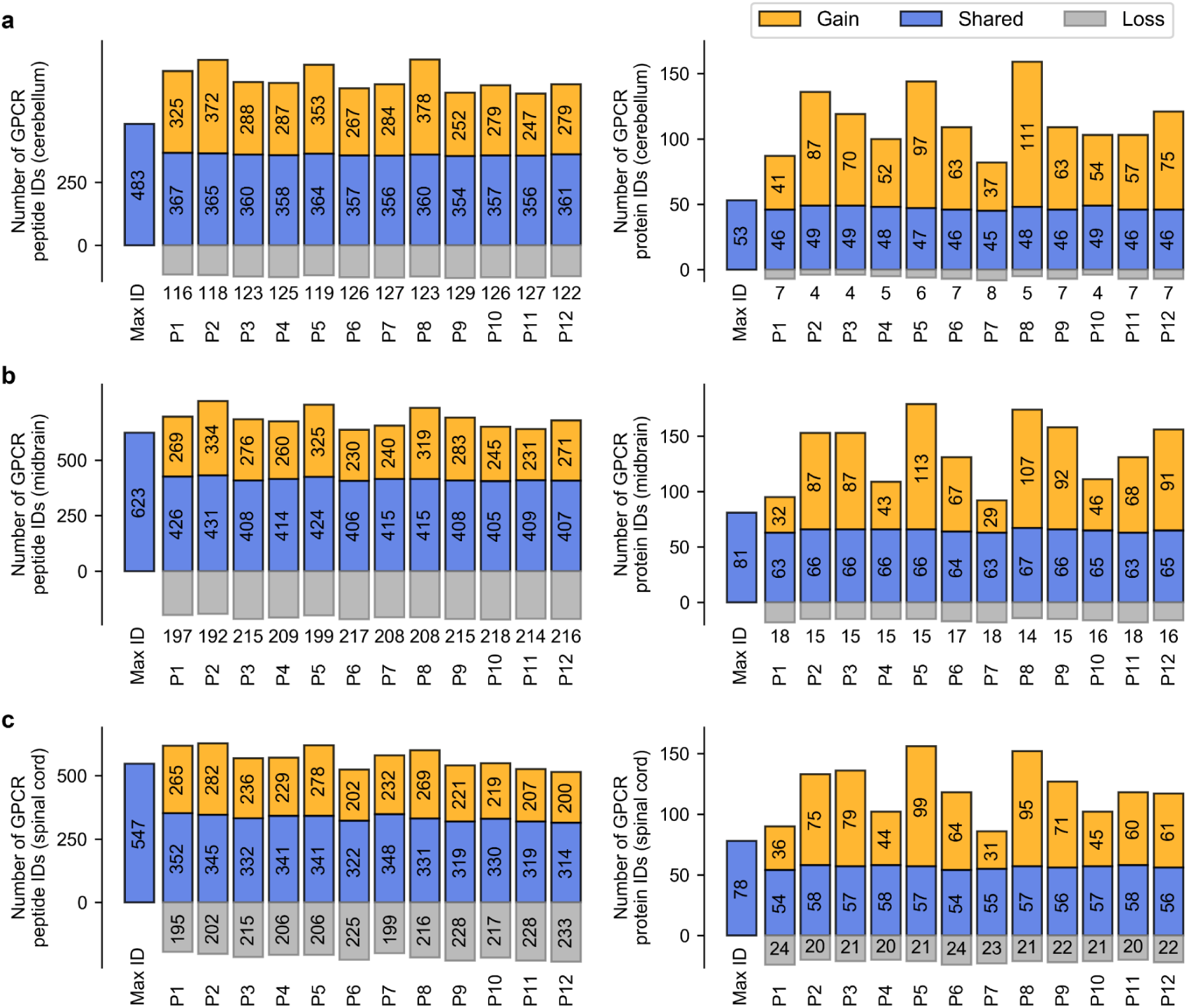
Comparison of GPCR identifications (IDs) in the max ID list of each brain region with those obtained using different hybrid libraries. Results are shown for the comparison of GPCR peptide IDs (left) and protein IDs (right) in cerebellum (a), midbrain (b), and spinal cord (c). Relative to the max ID list generated from DIA and DDA experiments (shown on the left), the proportion of shared IDs for each hybrid library is shown in blue; gained IDs are in orange; lost IDs are in grey. The number of protein IDs in each fraction is annotated.

**Supplementary Figure 8.**
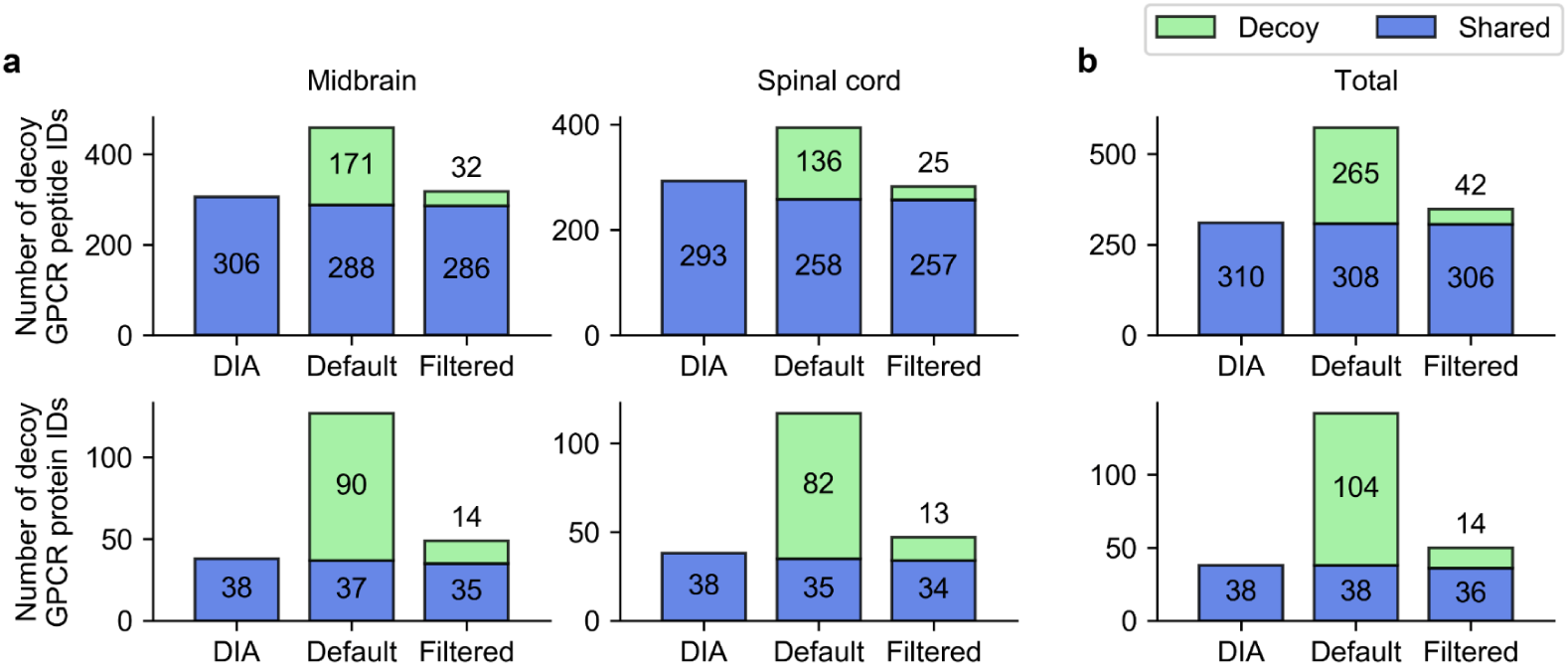
Data filtration removed a large portion of decoy GPCR peptides and proteins in two other brain regions. (a) Numbers of decoy GPCR peptide IDs (upper) and protein IDs (lower) in midbrain or spinal cord using the decoy hybrid library with default Spectronaut parameters (middle) or after data filtration with Cscore >0.9 (right). Relative to the protein/peptide IDs with the initial DIA library (shown on the left), the proportion of shared IDs is shown in blue and decoy IDs in green. (b) Numbers of total decoy GPCR peptide IDs (upper) and protein IDs (lower) in three brain regions using the decoy hybrid library with default Spectronaut parameters (middle) or after data filtration with Cscore >0.9 (right).

**Supplementary Figure 9.**
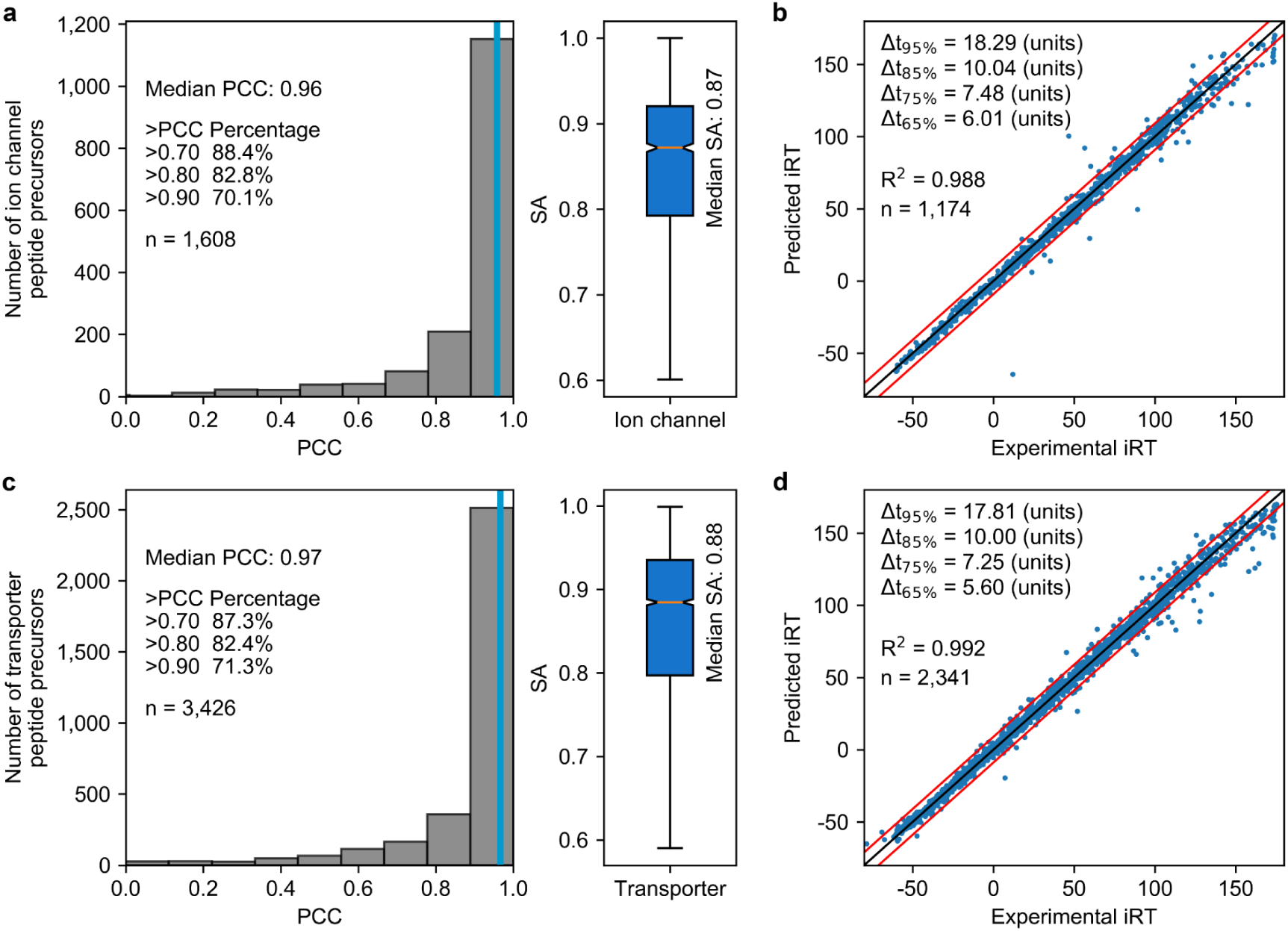
Testing re-trained models of pDeep and DeepRT in prediction for members in ion channel and transporter families. (a) Evaluation of the re-trained pDeep model based on distribution of PCC (left) and SA (right) calculated for ion channel peptides. (b) Evaluation of the re-trained DeepRT model based on correlation of predicted and experimental iRT values for ion channel peptides. (c) Evaluation of the re-trained pDeep model based on distribution of PCC (left) and SA (right) calculated for transporter peptides. (d) Evaluation of the re-trained DeepRT model based on correlation of predicted and experimental iRT values for transporter peptides. In each panel, n is the number of peptide precursors or peptides in the test set. In b and d, correlation coefficient (R^2^) is indicated, and red lines mark the iRT window required to encompass 95% of all peptides (Δ_t95%_) around the diagonal (black line).

**Supplementary Figure 10.**
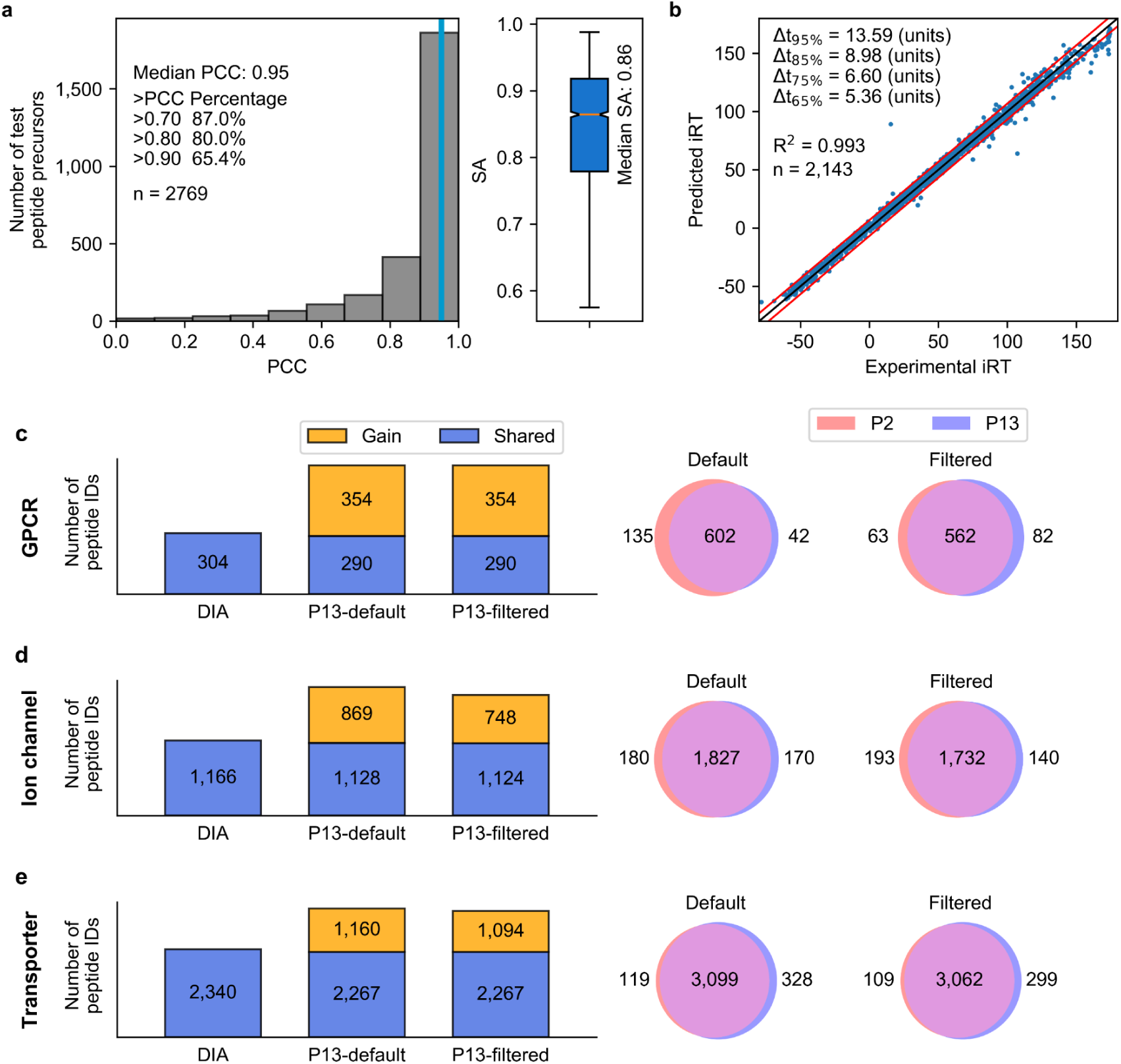
Building generic models that can predict for all three transmembrane protein families. (a) Evaluation of the generic pDeep model based on distribution of PCC (left) and SA (right) calculated for the test set. (b) Evaluation of the generic DeepRT model based on correlation of predicted and experimental iRT values for the test set. 90% of randomly selected precursors from the initial DIA library were used to re-train two models, and the remaining 10% precursors served as the test set. (c-e) Comparison of GPCR, ion channel and transporter peptide identifications (IDs) in cerebellum with the initial DIA library (DIA), or the new library P13 generated with the generic models. Shown left are numbers of peptide IDs using library P13 with default Spectronaut parameters or after data filtration (Cscore >0.9), and right are overlapping or distinct peptide IDs with default Spectronaut parameters or after data filtration.

**Supplementary Figure 11.**
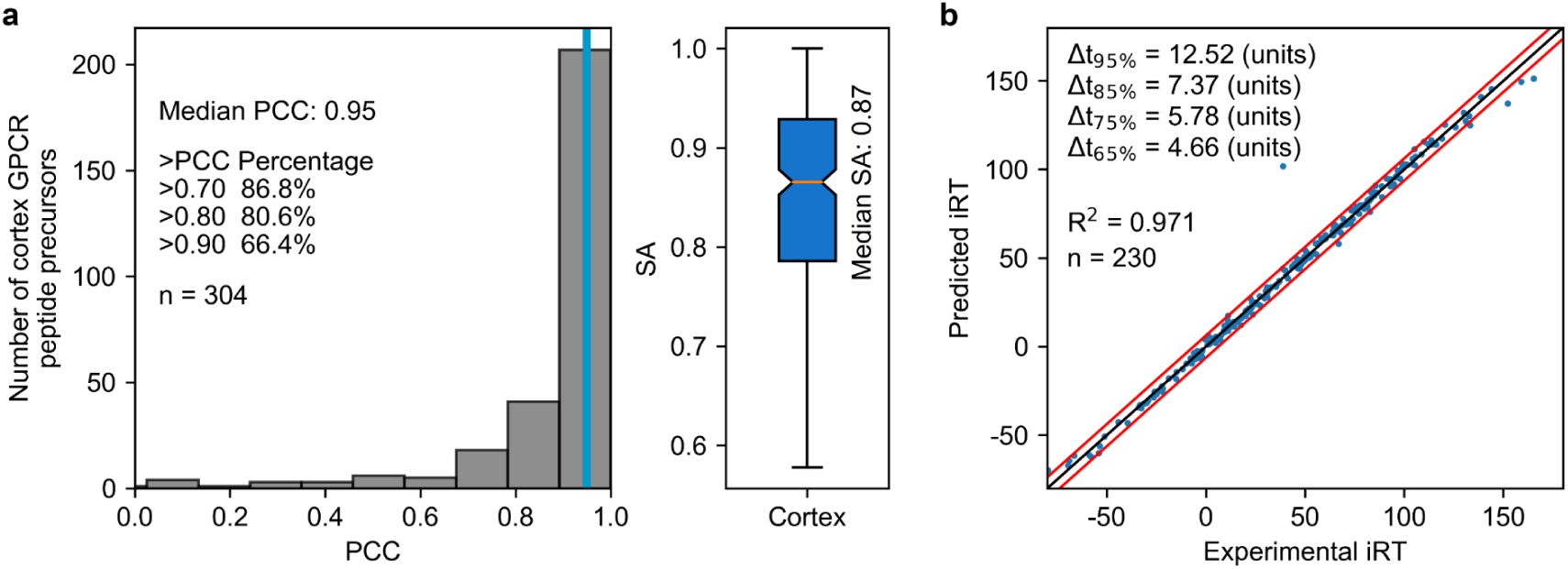
Testing models of pDeep and DeepRT re-trained with the published barrel cortex DIA dataset. Two models were used to predict fragmentation patterns and iRT values for mouse GPCR peptides in the test set. (a) Evaluation of the re-trained pDeep model based on distribution of PCC (left) and SA (right) calculated for GPCR peptide precursors. (b) Evaluation of the re-trained DeepRT model based on correlation of predicted and experimental iRT values for GPCR peptides. Correlation coefficient (R^2^) is indicated, and red lines mark the iRT window required to encompass 95% of all peptides (Δ_t95%_) around the diagonal (black line). In both panels, n is the number of peptide precursors or peptides in the test set.

**Supplementary Figure 12.**
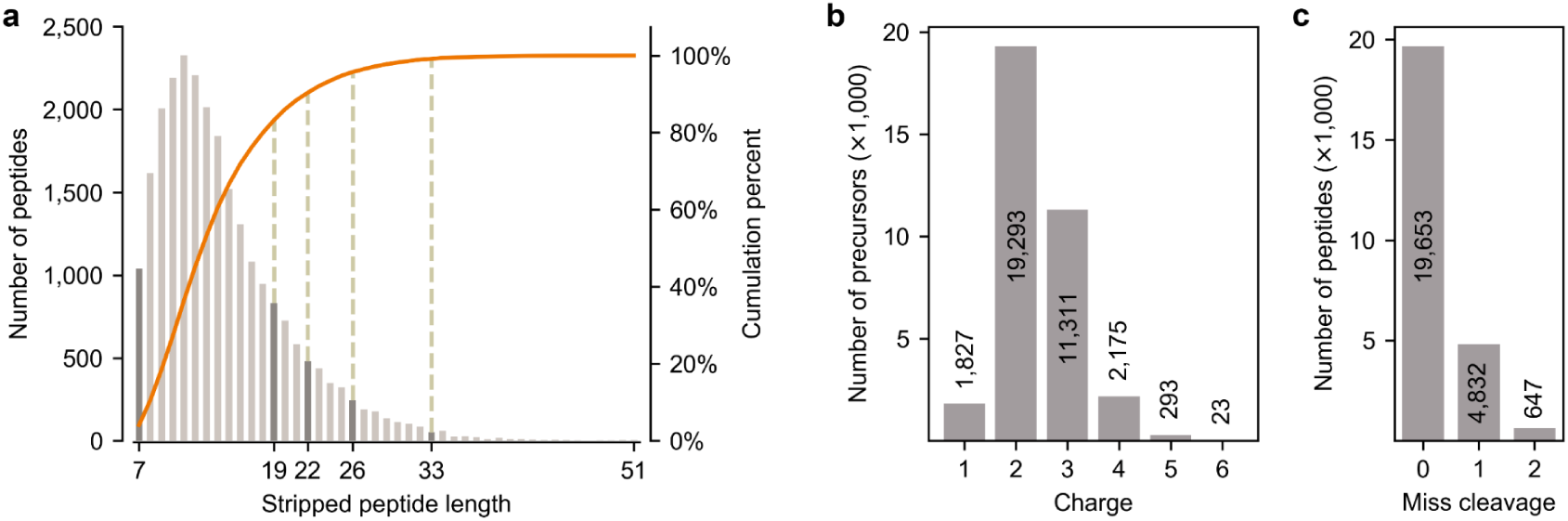
Properties of stripped peptides or precursors in the initial DIA library. (a) Length distribution of stripped peptides. Orange line associated with the right y-axis indicates the cumulated percentage of peptides of increased lengths. Above 95% of all peptides in the library have 7 to 26 residues, and above 99% of them have 7 to 33 residues. (b) Charge distribution of peptide precursors. Above 99% of all peptides in the library have 1 to 4 charges. (c) Distribution of missed cleavage number of stripped peptides.

